# *JAK2*^V617F^ Myeloproliferative Neoplasms Support Parallel Evolution of Independent Leukemic Clones

**DOI:** 10.1101/2025.09.23.678057

**Authors:** Tyler M. Parsons, Aishwarya Krishnan, Infencia Xavier Raj, Andrew L. Young, David R. O’Leary, Jason Arand, Maggie Cox, Stephen T. Oh, Grant A. Challen

**Affiliations:** Division of Oncology, Department of Medicine, Washington University School of Medicine; St. Louis, MO, USA, 63110; Division of Hematology, Department of Medicine, Washington University School of Medicine; St. Louis, MO, USA, 63110

## Abstract

Myeloproliferative neoplasms (MPNs) are hematological diseases predominantly driven by the *JAK2*^V617F^ mutation. Progression from chronic-phase MPN to secondary acute myeloid leukemia (sAML) is a severe complication that dramatically worsens disease prognosis. While progression to sAML is classically linked to MPN clones acquiring additional mutations, the absence of *JAK2*^V617F^ in some cases of post-MPN sAML cases suggests alternative mechanisms of transformation. Utilizing patient samples and *in vivo* modeling, we establish that leukemic clones can emerge independently of *JAK2*-mutant cells and undergo positive selection in the pro-inflammatory MPN environment, leading to parallel disease evolution. Genetic and pharmacological inhibition of IL-12 and TNFα mitigates this competitive advantage. Our data establish a new paradigm and show that disease progression in MPN can arise from parallel acute myeloid leukemia (pAML) clones.

## Main Text

Myeloproliferative neoplasms (MPNs) are a group of chronic hematological diseases driven by the acquisition of somatic mutations, predominantly *JAK2*^V617F^, in hematopoietic stem cells (HSCs)(*1–3*). MPNs are characterized by the aberrant and unregulated proliferation of one or more myeloid lineages, resulting in the overproduction of mature hematopoietic cells in the bone marrow (BM) and peripheral blood (PB)(*4–6*). This sustained overproduction manifests as polycythemia vera (PV; excess erythrocytes), essential thrombocythemia (ET; excess platelets), or myelofibrosis (MF; BM fibrosis). Consequently, patients face increased risks of blood viscosity and clotting, organ enlargement, joint pain and swelling, BM scarring, and are at-risk for progression to more aggressive disease states. The transformation of chronic-phase MPN to secondary acute myeloid leukemia (sAML) is a severe complication traditionally linked to the acquisition of additional mutations in the MPN driver clone in genes encoding epigenetic regulators (e.g. *TET2, DNMT3A, ASXL1)*, tumor suppressors (e.g. *TP53, JARID2)* and signaling molecules (e.g. *N*/*K*-*RAS)*(*7–10*). However, the *JAK2*^V617F^ driver mutation is occasionally absent in sAML that transforms from antecedent *JAK2*-mutant MPNs, which suggests alternative mechanisms of disease evolution(*11*). Proposed explanations for the absence of the *JAK2*^V617F^ mutation at the sAML stage include somatic reversion and loss of heterozygosity, but an underexplored possibility is that sAML originates from independent clones that evolve in parallel to the MPN during the natural history of clonal hematopoiesis (CH)(*12, 13*).

Here, we establish that pre-leukemic clones can arise independently of the MPN and outcompete the *JAK2*^V617F^-mutant cells to manifest disease progression, challenging the traditional dogma of post-MPN sAML evolution. We leveraged a series of primary patient samples **(Table S1)** and pre-clinical models to investigate the growth and evolution of parallel clones in an MPN background. We demonstrate that independent clones carrying *TET2* and *TP53* mutations are positively selected by MPN-derived pro-inflammatory cytokines such as IL-12 and TNFα. Importantly, we demonstrate these mechanisms are amenable to targeted therapy as genetic and pharmacological inhibition of IL-12- and TNFα-signaling ameliorates the competitive advantage of *TET2*-mutant cells in the presence of *JAK2*^V617F^-mutant MPN. Collectively, these data demonstrate that *JAK2*^V617F^-mutant cells condition an environment that confers a selective advantage for the parallel growth of independent clones in the background of existing MPN. In such cases, the more accurate diagnosis is two independent diseases evolving in the same patient – the primary MPN and a parallel acute myeloid leukemia (pAML). Improved understanding of the phylogeny of MPN disease evolution may offer new clinical opportunities for these patients who currently have very limited treatment options.

### Leukemic Clones Can Arise Independently and Undergo Positive Selection in a Background of MPN

To investigate the clonal trajectory post-MPN sAML, we performed single cell genomic analysis for three paired patient samples (2 MF; 1 PV) who progressed from a *JAK2*^V617F^-mutant MPN to a *JAK2*-negative sAML as defined by clinical sequencing **(Fig. S1A)** . Single-cell sequencing data from purified CD34^+^ cells for two patients (MF UPN:950899; PV UPN:374024) demonstrated that mutations detected at the sAML stage were not detected within the *JAK2*-mutant clones at the MPN stage (**Fig. 1A**). Despite clinical genomics defining MF UPN:638574 as *JAK2*-negative sAML, single cell analysis of purified CD34^+^ cells clearly showed the leukemic mutations were acquired in the founding MPN clone **(Fig. 1A)**. This was confirmed by droplet digital PCR (ddPCR) wherein the *JAK2*^V617F^ mutation was detected at both the MPN and the sAML stages **(Fig. S1B)**. While these data show there are multiple routes to transformation of post-MPN sAML, the single cell genomics clearly show that the leukemia-initiating mutations are not always present within the *JAK2*-mutant clones and can arise independently and outcompete the MPN cells to drive transformation.

**Fig. 1.**
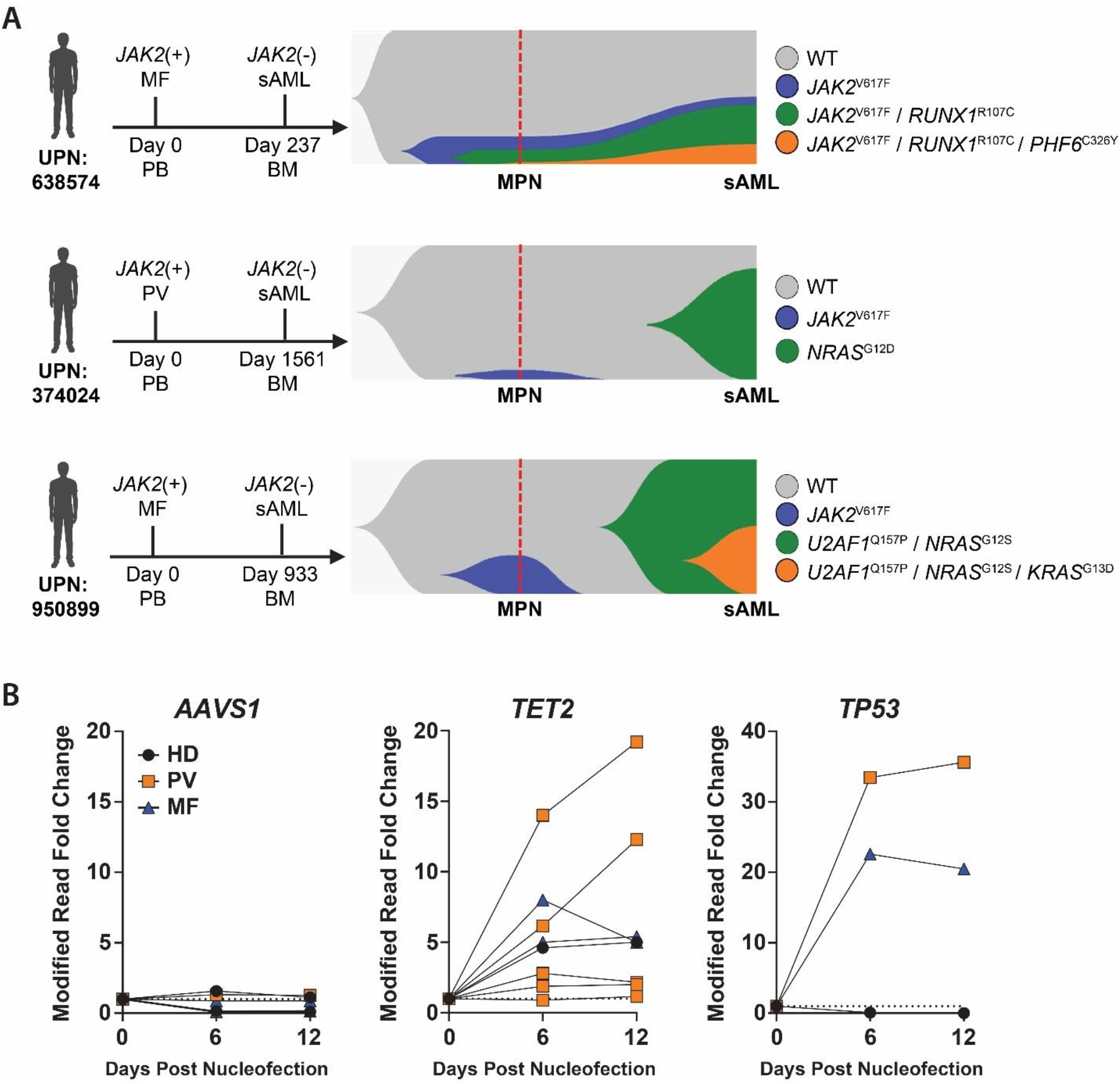
Leukemic Clones Can Arise Independently and Undergo Positive Selection in a Background of MPN. **(A)** Clonal hierarchy visualization of three paired (MPN and sAML) patient samples resolved by single-cell genomic sequencing with the x-axis expressed as time and the y-axis displaying proportionate prevalence of each clone within the population. **(B)** *ex vivo* competition assay showing VAF of engineered *AAVS1, TET2*, and *TP53* mutations in cord blood CD34^+^ cells in co-culture with CD34^+^ cells from *JAK2*^V617F^ PV or MF patients or healthy donor (HD) BM.

Given this finding, we developed an *ex vivo* competition system to quantify the proliferation of independent clones in the presence of MPN cells utilizing CRISPR/Cas9 editing. 1×10^5^ CD34^+^ cells from *JAK2*-mutant MPN patients or healthy donor (HD) BM were co-cultured for 12-days with 2×10^4^ cord blood (CB)-derived CD34^+^ cells nucleofected with gRNAs targeting *AAVS1, TET2* or *TP53*. CRISPR edited alleles were tracked via targeted genomic sequencing at 6-day intervals and compared to initial values 24-hours post-nucleofection. In the presence of MPN patient CD34^+^ cells, *TET2*- and *TP53*-mutant clones expanded significantly more compared to co-culture of the same clones with HD BM control CD34^+^ cells **(Fig. 1B)**. Interestingly, PV patient cells supported the growth of both *TET2*- and *TP53*-mutant cells moreso than MF patient cells, a finding consistent with clinical reports that the *JAK2*-mutant MPN to *JAK2* wild-type sAML trajectory is more common in PV patients(*11, 13*).

### *JAK2*^V617F^-Mutant MPN Accelerates Expansion of Independent *TET2*- and *TP53*-Mutant Clones

To explore competition dynamics between MPN cells and independent clones, we leveraged our MPN patient derived xenograft (PDX) system(*14*). *TET2* and *TP53* mutations were engineered into CB-derived CD34^+^ cells using CRISPR/Cas9 homology-directed repair (HDR) to introduce *TET2*^1216*^ and *TP53*^R248Q^ knock-in (KI) mutations into CB-derived CD34^+^ cells **(Fig. S2)**. These mutations were chosen from the limited literature describing *TET2* and *TP53* mutations as the most common variants in *JAK2*-negative post-MPN sAML(*15, 16*). A single-stranded oligo donor nucleotide (ssODN) was designed to introduce a silent mutation in the inert *AAVS1* locus to serve as a trackable negative control genetic barcode. As most *TET2* and *TP53* mutations in myeloid malignancies lead to loss of function(*17–22*), indels resulting from CRISPR/Cas9 cutting but failed ssODN directed repair were additionally tracked. These edits should also provide the cells with a competitive advantage and increase the clonal complexity within a given experiment. PDX models were established by co-transplanting 2.0×10^4^ cells from each of the *AAVS1, TET2* and *TP53* nucleofected populations into NSGS immunodeficient mice with 1.0×10^5^ CD34^+^ cells derived from either HD BM confirmed to be *JAK2*^V617F^-negative by ddPCR (control; n=4), *JAK2*-mutant MF (n=4), or *JAK2*-mutant PV (n=4) patients **(Table S1)**. Flow cytometric analysis was performed to confirm PB lineage reconstitution **(Fig. S3)**, and PDX models supported robust engraftment of human cells (**Fig. 2A**) in the BM (**Fig. 2B**). Notably, mice engrafted with PV patient cells supported significant expansion of human HSCs (mCD45-hCD45+ LINEAGE-CD34+ CD38-CD45RA-CD90+) in the BM (**Fig. 2C**). PDXs recapitulated classical phenotypes of the patients from which they were derived (WBC, HCT, splenomegaly; **Fig. 2D-F)** and served as a relevant platform to observe how different MPN subtypes support the expansion of independent *TET2*- and *TP53*-mutant clones. Targeted genomic sequencing was performed on hCD45+ cells isolated from the BM of recipient mice at the conclusion of each independent transplant (16-weeks) to quantify gRNA-mediated variant allele fraction (VAF). These data were compared to pre-transplant values to analyze the growth dynamics of parallel clones isolated from a *JAK2*^V617F^-mutant MPN-derived PDX host in comparison to the same pool of edited cells in a HD control host. MF and PV patient cells supported the growth of *TET2-* and *TP53-*mutant clones significantly more than HD control BM across four separate experiments established with independent starting MPN patient material **(Fig. 2G)**. Notably, cohorts established from PV patient cells exhibited the highest expansion of independent clones, particularly for *TET2*-mutant cells, mirroring results from the *ex vivo* experiment. These data highlight the robust expansion capacity of *TET2*- and *TP53*-mutant clones within *JAK2*^V617F^-mutant MPN environments *in vivo*.

**Fig. 2.**
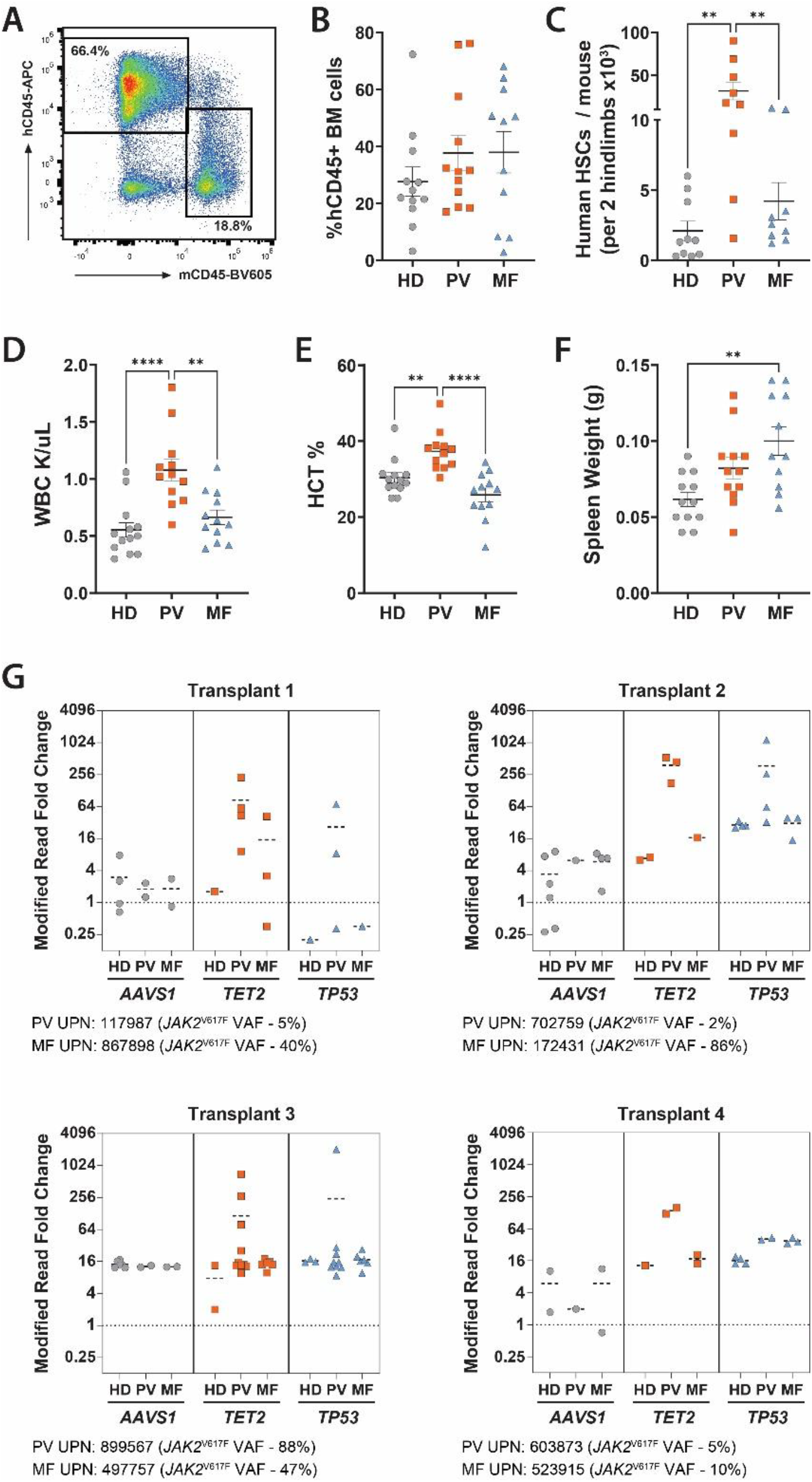
*JAK2*^V617F^-Mutant MPN Accelerates Expansion of Independent *TET2*- and *TP53*- Mutant Clones. Representative flow cytometry plot depicting human cell engraftment in BM of NSGS mice. Quantification of overall hCD45+ BM engraftment, **(C)** human HSC number, **(D)** white blood cell count, **(E)** hematocrit and **(F)** spleen weights compiled across PDX experiments. **(G)** Relative fold change of CRISPR modified reads in hCD45+ BM cells 16-weeks post-transplant compared to day 0 values of tracked *AAVS1, TET2*, and *TP53* engineered mutations in cord-blood CD34^+^ cells (n= 12 recipient mice sourced from 4 separate donors for HD, PV, and MF). ^**^*p* ≤0.01, ^****^*p* ≤0.0001

### Low Burden of *Jak2*^V617F^-Mutant Cells Specifically Supports Parallel Expansion of *Tet2*-Mutant Clones

With the finding that independent *TET2* and *TP53* clones display a significant growth advantage in a *JAK2*^V617F^-mutant environment, we established murine chimera models to provide a platform for mechanistic studies. These systems allow for precise control of the burden of each mutant cell population at the start of each transplant, facilitating a more detailed study of clone behavior and interaction than afforded by PDX models. We utilized mice with inducible expression of *Jak2*^V617F^ in the hematopoietic system (Vav-Cre;*Jak2*^V617F/+^ = “*Jak2*^V617F^”), which generate a PV-like phenotype, or wild-type (WT) control mice as a host background and *Tet2* heterozygous loss-of-function (Vav-Cre;*Tet2*^fl/+^ = “*Tet2*^Δ/+^”) or *Tp53* heterozygous knock-in *(Tp53*^R172H/+^) mice as the competing test populations. 2.5×10^6^ BM cells from either *Jak2*^V617F^ or WT control mice (CD45.2) were mixed with 5×10^5^ BM cells from either *Tet2*^Δ/+^, *Tp53*^R172H/+^, or WT test BM (CD45.1/2) and transplanted into lethally irradiated recipients (CD45.1) to produce a starting fraction of “test” BM of approximately 15%.

Flow cytometric analysis was performed to evaluate PB lineage reconstitution **(Fig. S4)**. By 16-weeks post-transplant, *Jak2*^V617F^-mutant host cells significantly supported expansion of both *Tet2*^Δ/+^ and *Tp53*^R172H/+^ test populations in the PB compared to a WT control host background (**Fig. 3A-B**). However, the WT test population also exhibited an engraftment increase in the presence of *Jak2*^V617F^-mutant cells **(Fig. 3B)**. This engraftment advantage was largely restricted to peripheral lymphoid cells, consistent with prior reports that *JAK2*^V617F^ mutations confer lymphoid deficiency. Within the PB myeloid compartment in a *Jak2*^V617F^-mutant background, the WT test population exhibited significantly reduced chimerism compared to *Tet2*^Δ/+^ and *Tp53*^R172H/+^ test populations **(Fig. 3C-D)**. BM analysis 18-weeks post-transplant revealed a similar competitive advantage of *Tet2*^Δ/+^ and *Tp53*^R172H/+^ cells in a *Jak2*^V617F^-mutant background **(Fig. 3E)**. Interestingly, the effect was most pronounced in the hematopoietic stem/progenitor cell (HSPC; c-Kit+ Sca-1+ Lineage-“KSL”) compartment rather than the most primitive long-term hematopoietic stem cells (HSCs; KSL CD48-CD150+)**(Fig. 3F-G)**.

**Fig. 3.**
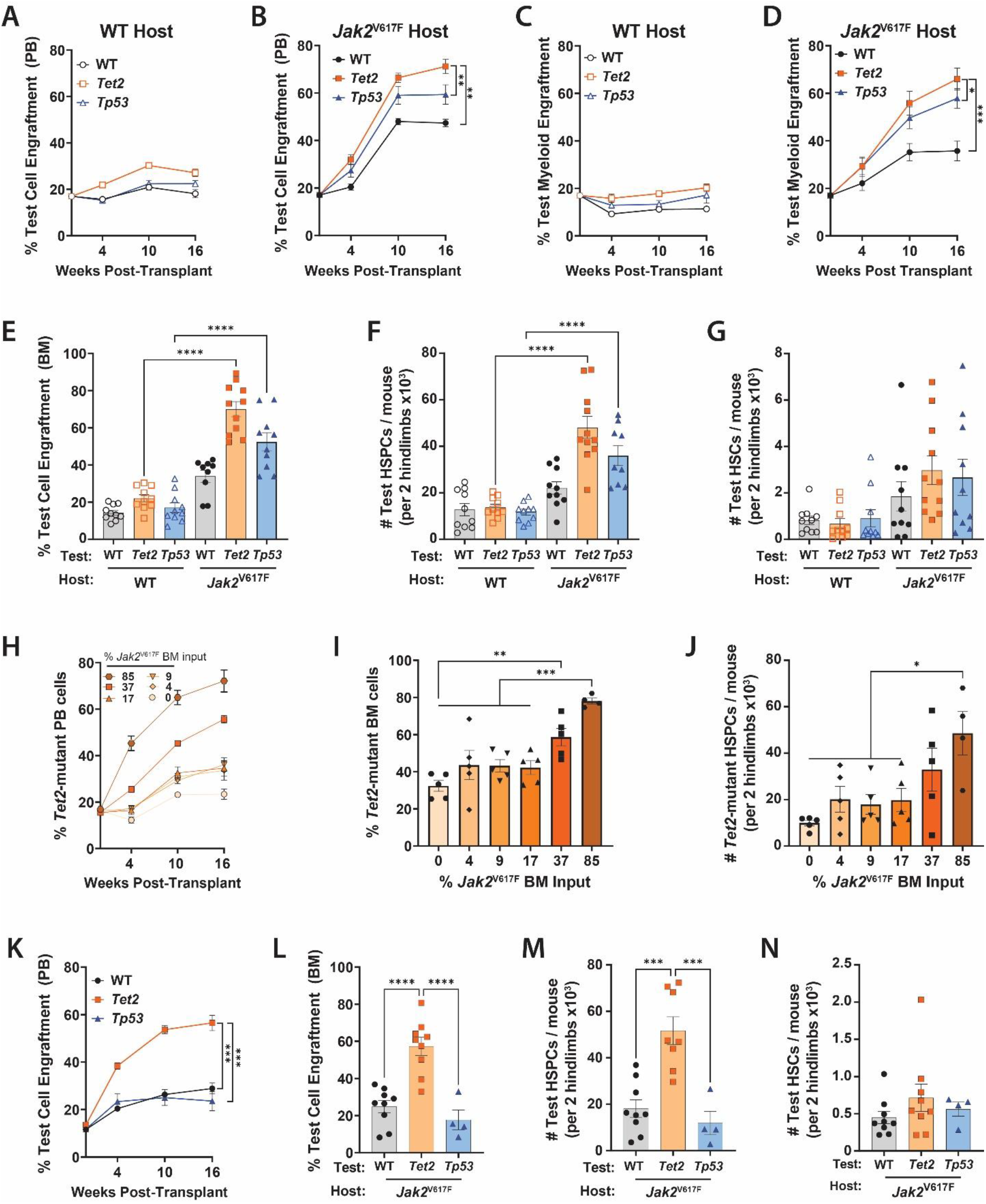
Low Burden of *Jak2*^V617F^-Mutant Cells Specifically Supports Parallel Expansion of *Tet2*-Mutant Clones. **(A, B)** Peripheral blood engraftment of WT control, *Tet2*^Δ/+^ and *Tp53*^R172H/+^ test cells in (A) WT or (B) a *Jak2*^V617F^ host BM. **(C, D)** PB myeloid (Gr-1^+^ Cd11b^+^) engraftment of WT control, *Tet2*^Δ/+^ and *Tp53*^R172H/+^ test cells in (C) WT or (D) *Jak2*^V617F^ host BM. **(E-G)** Quantification of WT, *Tet2*^Δ/+^, *Tp53*^R172H/+^ test cell (E) BM engraftment, (F) HSPC (Lineage^-^ Sca-1^+^ c-Kit^+^) number and (G) HSC (Lineage^-^ Sca-1^+^ c-Kit^+^ CD48^-^ CD150^+^) number in the BM of recipient mice 18-weeks post-transplant (n =10 / group). **(H-J)** Quantification of WT control and *Tet2*^Δ/+^ cell (H) PB engraftment, (I) BM engraftment and (J) HSPC numbers in recipient mice transplanted with indicated fractions of *Jak2*^V617F^ BM input cells (n = 5 / group). **(K-N)** Quantification of WT control, *Tet2*^Δ/+^, *Tp53*^R172H/+^ test cell (K) PB engraftment, (L) BM engraftment, (M) HSPC numbers and (N) HSC numbers in recipient mice transplanted with 50% *Jak2*^V617F^ BM input (n= 5-9 / group). ^*^*p* ≤0.05, ^**^*p* ≤0.01, ^***^*p* ≤0.001, ^****^*p* ≤0.0001

As *Tet2*-mutant clones demonstrated an enhanced growth advantage in a *Jak2*^V617F^-environment compared to *Tp53*-mutant clones in both PDX and murine chimeras, we aimed to determine the minimum *Jak2*^V617F^-mutant cell burden required to support parallel *Tet2*-mutant clone expansion. A titration experiment was performed wherein BM cells from *Jak2*^V617F^ mice were diluted at predetermined ratios with WT BM while maintaining a fixed *Tet2*^Δ/+^ cell input (15%). There was a threshold effect between starting *Jak2*^V617F^ cell input and the conferred competitive advantage of *Tet2*^Δ/+^ cells with a ∼35% *Jak2*^V617F^ mutant cell burden being the threshold needed to support robust expansion of *Tet2*^Δ/+^ cells **(Fig. 3H-J)**.

Based on these data, we established chimeras with 50% *Jak2*^V617F^-mutant BM, 35% WT support BM, and 15% *Tet2*^Δ/+^, *Tp53*^R172H/+^ or WT test BM to determine if this was also sufficient to induce outgrowth of parallel *Tp53*-mutant cells. As anticipated, a 50% starting *Jak2*-input supported positive selection of *Tet2*^Δ/+^ cells in the PB, BM, and HSPC compartments. However, this *Jak2*^V617F^-mutant burden did not support engraftment of *Tp53*^R172H/+^ or WT cells unlike the higher (85%) *Jak2*^V617F^-mutant cell input **(Fig. 3K-M)**. These findings highlight a specific interaction between *Jak2*^V617F^-mutant MPN cells and independent *Tet2*-mutant clones.

### IL-12 and TNFα Drive Expansion of *Tet2*-Mutant Clones in a *Jak2*^V617F^-Mutant Environment

As our data show a particularly strong selection of independent *Tet2*-mutant clones in an MPN background, we focused mechanistic studies to define this relationship. As the selective advantage for *Tet2*^Δ/+^ cells in and MPN background was most evident at the level of HSPCs, gene expression profiling was performed by RNA-sequencing analysis of *Tet2*^Δ/+^ HSPCs (CD45.1/2+ c-Kit+ Sca-1+ Lineage-) isolated from either WT or *Jak2*^V617F^-mutant hosts 18-weeks post-transplant. Analysis of differentially expressed genes (DEGs; **Fig. 4A**) demonstrated that *Tet2*^Δ/+^ HSPCs from a *Jak2*^V617F^-mutant environment exhibited a pronounced proliferation bias, exemplified by upregulation of *Mki67*, and increased expression of pro-myeloid differentiation genes such as *Elane, Mpo*, and *Cebpe*. The inflammatory response was also heightened with elevated levels of *S100a8* and *S100a9* which function as key promoters of myeloid inflammation and are routinely upregulated in HSPCs in the contexts of myeloid skewing and chronic inflammatory states(*23*). This myeloid differentiation bias occurred at the expense of lymphoid priming marked by decreased expression of *Dntt* and *Lck*, essential genes for lymphoid lineage commitment. Genes classically associated with HSC identity, such as *Hoxa10, Fgd5, Vldlr, Hlf*, and *Mecom*, were also downregulated in *Tet2*^Δ/+^ HSPCs from a *Jak2*^V617F^-mutant environment **(Fig. 4B)**, suggesting these cells were being pushed towards proliferation at the expense of self-renewal. Collectively, these findings indicate that a *Jak2*^V617F^-mutant environment drives *Tet2*-mutant HSPCs toward a proliferative, pro-myeloid, inflammatory phenotype. We next interrogated whether the gene expression changes observed in *Tet2*^Δ/+^ HSPCs from a *Jak2*^V617F^-mutant environment were due to specific effects on *Tet2*^Δ/+^ cells and not a general effect for any HSPCs exposed to MPN cells by RNA-seq comparison of *Tet2*^Δ/+^ and WT HSPCs isolated from a *Jak2*^V617F^-mutant environment. Differential expression analysis confirmed that the transcriptomic alterations previously identified in *Tet2*^Δ/+^ HSPCs remained dysregulated in *Tet2*^Δ/+^ HSPCs compared to WT HSPCs from a *Jak2*^V617F^-mutant environment **(Fig. S5)**. These findings demonstrate that the identified gene dysregulation is unique to *Tet2*-mutant cells within a *Jak2*^V617F^-mutant milieu and not a generalized feature of HSPCs exposed to MPN cells.

**Fig. 4.**
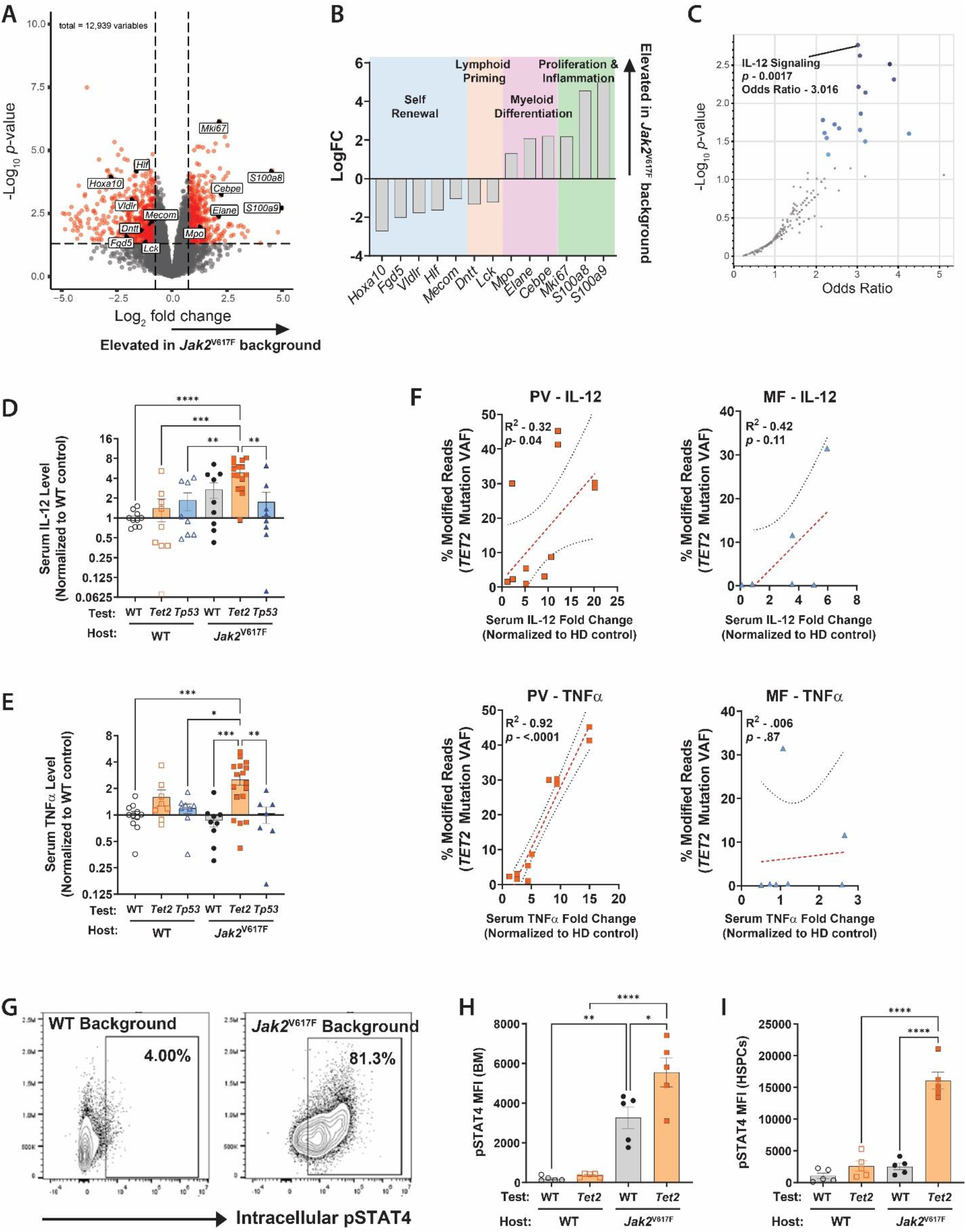
IL-12 and TNFα Drive Expansion of *Tet2*-Mutant Clones in a *Jak2*^V617F^-Mutant Environment. **(A)** Volcano plot showing differentially expressed genes between *Tet2*^Δ/+^ HSPCs from a *Jak2*^V617F^ environment compared to *Tet2*^Δ/+^ HSPCs from a WT environment. **(B)** LogFC values for *Tet2*^Δ/+^ HSPCs DEGs from indicated pathways. **(C)** Volcano plot of gene set over-representation analysis (ORA) identifying IL-12 signaling as the most significantly enriched pathway in *Tet2*^Δ/+^ HSPCs from a *Jak2*^V617F^ environment. **(D-E)** Serum cytokine levels (normalized to WT control chimeras) of (D) IL-12 and (E) TNFα from murine mixed chimeras (n= 9-15 / group). (**F**) Serum cytokine fold change (normalized to HD control mice) for PV and MF PDXs plotted against relative increase in VAF of individually tracked *TET2* mutations engineered into cord blood CD34^+^ cells (n= 12 recipient mice sourced from 4 separate donors for each HD, PV, and MF). **(G)** Representative flow cytometry plots showing intracellular pSTAT4 staining in *Tet2*^Δ/+^ BM cells in WT or *Jak2*^V617F^ backgrounds. **(H-I)** Median fluorescent intensity (MFI) of intracellular pSTAT4 levels in (H) whole BM and (I) HSPCs (n = 5 / group). ^*^*p* ≤0.05, ^**^*p* ≤0.01, ^***^*p* ≤0.001, ^****^*p* ≤0.0001

To discern the most dysregulated signaling pathways that may be driving the observed transcriptional changes, over-representation analysis (ORA) was performed on the identified DEGs. ORA identified IL-12-mediated signaling as the most significantly dysregulated cancer-related pathway between *Tet2*-mutant HSPCs isolated from a *Jak2*^V617F^-mutant background compared to from a WT background **(Fig. 4C)**. To further classify over-represented pathways, DEGs were quantitatively scored based on how frequently they appear in a cancer-related pathway relative to their frequency across all cancer-related pathways and used to weight gene set terms by their relevance across the dataset. Uniform manifold approximation and projection (UMAP) was then applied for dimensionality reduction to visualize relationships between enriched pathways. Again, among the identified clusters, IL-12-mediated signaling emerged as the most highly enriched pathway. Adjacent points in the UMAP space, inferring biological relevance of over-represented gene sets, included TNF receptor signaling and IL-12 signaling mediated by STAT4, highlighting these related signaling pathways as potential selection factors for *Tet2*^Δ/+^ HSPC expansion in a *Jak2*^V617F^-mutant environment **(Fig. S6)**.

It is well-documented that MPN cells secrete high levels of many pro-inflammatory cytokines(*24– 27*). Moreover, we and others have shown that many common CH mutations impart growth advantages to the mutant clones under different conditions of inflammation(*28–32*). To determine if the observed gene expression changes in *Tet2*^Δ/+^ HSPCs in a *Jak2*^V617F^-mutant environment were associated with altered cytokine levels due to the MPN cells, global cytokine profiling of PB serum from murine chimera and PDX experiments was performed. The overlap of cytokines increased in MPN models compared to relevant control comparators revealed four candidates - IL-1, IL-12, IL-27, and TNFα **(Fig. S7A)**. We sought to determine if any of these cytokines might create a competitive advantage for *TET2*-mutant clones. CB-derived CD34^+^ cells were CRISPR/Cas9-engineered with *TET2* mutations and cultured with each candidate cytokine at concentrations ranging from 1-100 ng/mL for 6-days. TNFα and IL-12 were the most effective in accelerating *TET2*-mutant cell proliferation *in vitro* **(Fig. S7B)**. Moreover, from *in vivo* studies, increasing IL-12 and TNFα levels correlated with *TET2*-mutant clonal expansion in murine (**Fig. 4D,E**) and PDX transplants **(Fig. 4F)**. This effect was most pronounced in PDX mice established from PV patient cells **(Fig. 4F)**. To determine the source of the aberrant signaling, IL-12 and TNFα levels were measured in serum from transplant donor mice of each genotype. IL-12 levels were significantly elevated in *Jak2*^V617F^-mutant mice compared to WT and *Tet2*^Δ/+^ donor mice. Conversely, TNFα was significantly elevated in *Tet2*^Δ/+^ donor mice **(Fig. S8)**. Notably, TNFα has been reported to correlate with *TET2*-mutant CH and IL-12 is a known TNFα stimulator that is elevated in MPN patients(*33–38*). This suggests a model whereby IL-12 secreted by MPN cells induces TNFα over-production by *Tet2*-mutant cells to condition an environment that fosters their development.

We hypothesized that IL-12 secreted by MPN cells directly acts upon *TET2*-mutant cells to phosphorylate STAT4, a key mediator of IL-12 signaling(*39–41*). Intracellular flow cytometry analysis (**Fig. 4G**) revealed that pSTAT4 levels were elevated in *Tet2*^Δ/+^ BM cells isolated from a *Jak2*^V617F^-mutant background 10-weeks post-transplant compared to those isolated from a WT background or from WT cells from either background **(Fig. 4H)**. Strikingly, the pSTAT4 elevation was most significant in the *Tet2*^Δ/+^ HSPC population isolated from a *Jak2*^V617F^-mutant environment (**Fig. 4I**). This indicates a potential mechanistic link between IL-12 signaling and *Tet2*^Δ/+^ HSPC proliferation in a *Jak2*^V617F^-mutant context, suggesting that targeting this pathway could mitigate the competitive advantage in this genetic setting.

### Inhibition of Inflammatory Cytokines Mitigates the Competitive Advantage of *TET2*-Mutant Cells in a *Jak2*-Mutant Environment

With the identification of IL-12 and TNFα as potential selective pressures for independent *Tet2*-mutant clone expansion in parallel to MPN, we aimed to evaluate the functional consequences of disrupting these pathways. To enhance the clinical relevance, we replicated mutational burdens that more closely mirror those observed in MPN patients with input thresholds determined by the *Jak2*^V617F^ titration experiment. As such, chimeras utilized in functional studies were established by transplanting 1.5×10^6^ *Jak2*^V617F^ BM cells (CD45.2) with 4.5×10^5^ test BM (CD45.1/2) and 1.05×10^6^ WT support cells (CD45.1) into lethally irradiated recipients (CD45.1) to produce a starting fraction of test BM of approximately 15% in an environment composed of approximately 50% *Jak2*^V617F^-mutant host BM. First, to assess the implications of TNFα on the observed competitive advantage of *Tet2*-mutant clones, Vav-Cre;*Tet2*^fl/+^ mice were crossed with a TNFα-receptor genetic deletion model (p55/p75 germline knockout = “KO”) to establish mice wherein *Tet2*-mutant hematopoietic cells lack receptors for TNFα. To determine the effect of ameliorating TNFα signaling in *Tet2*-mutant cells, BM cells from either *Jak2*^V617F^ or WT mice were mixed with either *Tet2*^Δ/*+*^, *TNFαr*p55^-/-^p75^-/-^ (= “*TNFr*^KO^”), *Tet2*^Δ/+^/*TNFr*^KO^, or WT test BM and co-transplanted with WT support. Blood **(Fig. 5A-C)** and BM analysis **(Fig. 5D)** revealed the competitive advantage of *Tet2*-mutant cells in a *Jak2*^V617F^-mutant environment was significantly blunted by genetic deletion of TNFα receptors on the *Tet2*-mutant cells.

**Fig. 5.**
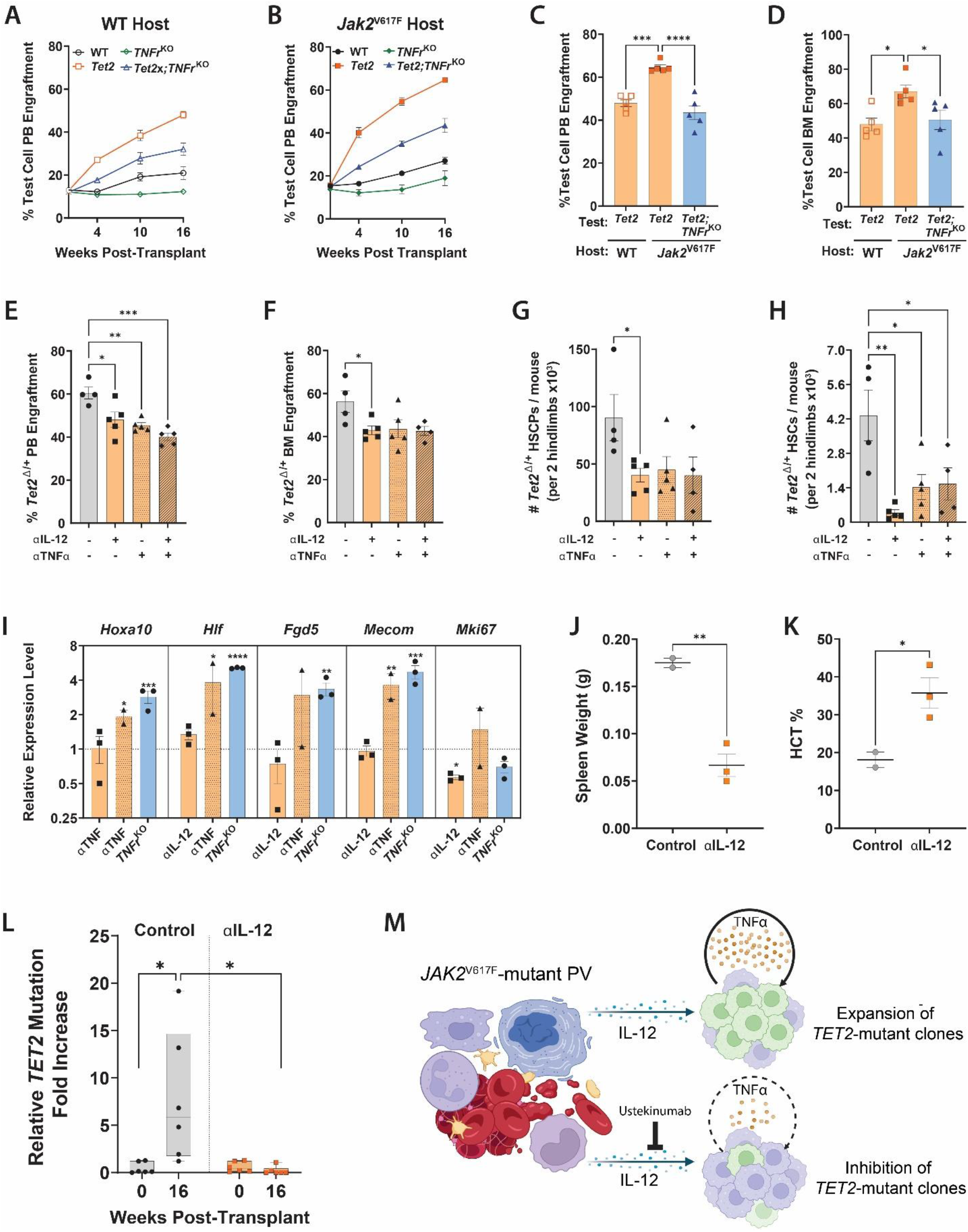
Inhibition of Inflammatory Cytokines Mitigates the Competitive Advantage of *TET2*-Mutant Cells in a *Jak2*-Mutant Environment. **(A, B)** Peripheral blood engraftment of WT control, *Tet2*^Δ/+^, *TNFr*^KO^ and *Tet2*^Δ/+^;*TNFr*^KO^, test cells in (A) WT or (B) a *Jak2*^V617F^ host BM (n= 5 / group). (**C**) 16-week peripheral blood engraftment of *Tet2*^Δ/+^ and *Tet2*^Δ/+^, *TNFr*^KO^ test cells in WT or *Jak2*^V617F^ host BM (n= 5 / group). (**D**) 18-week BM engraftment of *Tet2*^Δ/+^ and *Tet2*^Δ/+^, *TNFr*^KO^ test cells in WT or *Jak2*^V617F^ host BM (n= 5 / group). (**E-H**) *Tet2*^Δ/+^ test cell (E) 16-week PB engraftment, (F) 18-week BM engraftment, (G) HSPC number and (H) HSC number chimeric mice receiving neutralizing antibodies against IL-12 and/or TNFα (n= 4-5 / group). **(I)** qPCR gene expression analysis of *Tet2*^Δ/+^ HSPCs from indicated treatments of genetic models depicting levels of indicated genes (normalized to non-treated controls; n = 2-3 / group). **(J-L)** Analysis of (J) spleen weight, (K) hematocrit and (L) modified read fold change for CRISPR engineered *TET2* mutations in cord blood CD34^+^ cells in treatment-control and IL-12-neutralizer treated PDX mice co-transplanted with PV UPN:702759 CD34^+^ cells (n= 2-3 / group). **(M)** Schematic depicting mechanism of PB cells providing a selective advantage to independent *TET2*-mutant clones through cytokine support. ^*^*p* ≤0.05, ^**^*p* ≤0.01, ^***^*p* ≤0.001, ^****^*p* ≤0.0001

To evaluate this finding in a more translational system, we administered murine biosimilars of TNFα and IL-12 monoclonal neutralizing antibodies (adalimumab / ustekinumab respectively) into mouse chimeric models between weeks 4-10 post-transplant. Inhibition of TNFα and IL-12 dramatically reduced *Tet2*^Δ/+^ cell engraftment in a *Jak2*^V617F^-mutant background in the PB **(Fig. 5E)**, BM **(Fig. 5F)** and HSPC populations **(Fig. 5G)**. Notably, neutralization of IL-12 and TNFα suppressed *Tet2*^Δ/+^ cells down to the HSC level – an effect most significantly achieved by IL-12 neutralization **(Fig 5H)**. Gene expression analysis of *Tet2*^Δ/+^ HSPCs isolated from a *Jak2*^V617F^-mutant background at the end of the 18-week study from control and treated cohorts showed targeting the IL-12/TNFα axis normalized dysregulation in proliferation *(Mki67)* and self-renewal *(Hoxa10, Hlf, Fgd5*, and *Mecom)* pathways. Similar normalization of these abnormal gene expression programs was also observed in *Tet2*^Δ/+^ HSPCs genetically deficient for TNFα receptors **(Fig. 5I)**. Thus, genetic and pharmacological inhibition of aberrant MPN cytokine signaling abrogates the competitive advantage of *Tet2*^Δ/+^ HSPCs in the presence of *Jak2*^V617F^-mutant cells. To evaluate the effect of IL-12 neutralization in a human system, we utilized a *JAK2*^V617F^-mutant PV patient sample that previously demonstrated support of parallel *TET2*-mutant clones (UPN:702759). 1×10^5^ CD34^+^ PV patient cells were co-transplanted with 2.0×10^4^ CB-derived CD34^+^ cells harboring gRNA-mediated *TET2* mutations. Between weeks 10-16 post-transplant, a human IL-12 neutralizing agent was administered to the treatment group. Control mice exhibited signs of disease progression characterized by increased spleen weight and decreased hematocrit, while mice receiving the IL-12 neutralizing agent retained PV-like pathologies **(Fig. 5J-K)**. Strikingly, IL-12 neutralization was able to mitigate the competitive advantage of *TET2*-mutant clones in the presence of PV patient cells **(Fig. 5L)**. These findings reveal that targeting IL-12 could be a potential therapeutic approach for restricting *TET2*-mutant CH in the background of an existing *JAK2*^V617F^-mutant driven MPN to minimize future risk of disease progression to pAML **(Fig. 5M)**.

## Conclusion

These data establish that *JAK2*^V617F^-mutant cells potentiate parallel expansion of independent clones as a non-classical trajectory of disease progression in MPN. We show that MPN cells drive this parallel evolution through an IL-12/TNFα cytokine axis, which biases *Tet2*-mutant progenitors towards increased proliferation and myeloid differentiation. Genetic and pharmacological inhibition of IL-12 and TNFα resulted in both functional and molecular rescue of these phenotypes. Our findings provide crucial insights into how sAML which lacks the *JAK2*^V617F^ driver mutation can evolve from an antecedent *JAK2*^V617F^-mutant MPN. These results highlight a specific example of clonal dynamics and cell competition within a *JAK2*^V617F^-mutant context; however, it is important to acknowledge that clones with other CH mutations (e.g. *DNMT3A)* as well as MPNs driven by other mutations *(CALR, MPL)* may confer different patterns of clonal evolution. We aim to leverage these findings to enhance disease surveillance in MPN populations. With improved genomic monitoring, targeted interventional therapies to prevent the expansion of emerging pre-leukemic clones could reduce risk of parallel disease evolution in MPN, a significant therapeutic advantage for a substantial subset of patients.

## Acknowledgments

We thank all members of the Challen laboratory. We thank the Alvin J. Siteman Cancer Center at Washington University School of Medicine and Barnes-Jewish Hospital in St. Louis, MO. for the use of the Siteman Flow Cytometry Core facilities supported in part by an NCI Cancer Center Support Grant #P30CA091842. We thank Dr. Jorge Di Paola for generously providing TNFα-receptor germline knock-out mice. *Tp53*^R172H^ mice were originally developed by Dr. Guillermina Lozano (MD Anderson Cancer Center). This publication is solely the responsibility of the authors and does not necessarily represent the official views of the NIH. T.M.P is a Fellow of Blood Cancer United. A.K. was supported by the American Society of Hematology (ASH) Graduate Hematology Award. A.L.Y. was supported by the ASH Research Training Award for Fellows and Scholar Award, and the Edward P. Evans Foundation Young Investigator Award. I.X.R. was supported by NIH P30 CA091842.

## Author Contributions

Conceptualization: TMP, STO, GAC

Methodology: TMP, AK, IXR, ALY, GAC

Investigation: TMP, AK, IXR, ALY, DO, JA, MC, GAC

Visualization: TMP, IXR, ALY

Funding acquisition: GAC

Project administration: TMP, JA, GAC

Supervision: GAC

Writing – original draft: TMP, GAC

Writing – review and editing: TMP, GAC

## Data and materials availability

Data not available in the main text or the supplementary materials are available through request to the corresponding author. All raw read data (FASTQ files) for RNA sequencing are publicly available at the Gene Expression Omnibus database accession number GSE308233.

## Supplementary Materials

Materials and Methods

Figs. S1 to S8

Table S1 to S2

References (42-50)

Data S1

## References and Notes

1. R. Kralovics, F. Passamonti, A. S. Buser, S.-S. Teo, R. Tiedt, J. R. Passweg, A. Tichelli, M. Cazzola, R. C. Skoda, A Gain-of-Function Mutation of JAK2 in Myeloproliferative Disorders. N Engl J Med 352, 1779–1790 (2005).

2. E. J. Baxter, L. M. Scott, P. J. Campbell, C. East, N. Fourouclas, S. Swanton, G. S. Vassiliou, A. J. Bench, E. M. Boyd, N. Curtin, M. A. Scott, W. N. Erber, A. R. Green, Acquired mutation of the tyrosine kinase JAK2 in human myeloproliferative disorders. The Lancet 365, 1054–1061 (2005).

3. R. L. Levine, M. Wadleigh, J. Cools, B. L. Ebert, G. Wernig, B. J. P. Huntly, T. J. Boggon, I. Wlodarska, J. J. Clark, S. Moore, J. Adelsperger, S. Koo, J. C. Lee, S. Gabriel, T. Mercher, A. D’Andrea, S. Fröhling, K. Döhner, P. Marynen, P. Vandenberghe, R. A. Mesa, A. Tefferi, J. D. Griffin, M. J. Eck, W. R. Sellers, M. Meyerson, T. R. Golub, S. J. Lee, D. G. Gilliland, Activating mutation in the tyrosine kinase JAK2 in polycythemia vera, essential thrombocythemia, and myeloid metaplasia with myelofibrosis. Cancer Cell 7, 387–397 (2005).

4. S. Anand, F. Stedham, P. Beer, E. Gudgin, C. A. Ortmann, A. Bench, W. Erber, A. R. Green, B. J. P. Huntly, Effects of the JAK2 mutation on the hematopoietic stem and progenitor compartment in human myeloproliferative neoplasms. Blood 118, 177–181 (2011).

5. A. R. Moliterno, H. Kaizer, B. N. Reeves, JAK2V617F Allele Burden in Polycythemia Vera: Burden of Proof. Blood, blood.2022017697 (2023).

6. J. Grabek, J. Straube, M. Bywater, S. W. Lane, MPN: The Molecular Drivers of Disease Initiation, Progression and Transformation and their Effect on Treatment. Cells 9, 1901 (2020).

7. W. Walter, N. Nadarajah, S. Hutter, H. Müller, C. Haferlach, W. Kern, T. Haferlach, M. Meggendorfer, Characterization of myeloproliferative neoplasms based on genetics only and prognostication of transformation to blast phase. Leukemia 38, 2644–2652 (2024).

8. N. Maslah, L. Benajiba, S. Giraudier, J.-J. Kiladjian, B. Cassinat, Clonal architecture evolution in Myeloproliferative Neoplasms: from a driver mutation to a complex heterogeneous mutational and phenotypic landscape. Leukemia 37, 957–963 (2023).

9. O. Abdel-Wahab, T. Manshouri, J. Patel, K. Harris, J. Yao, C. Hedvat, A. Heguy, C. Bueso-Ramos, H. Kantarjian, R. L. Levine, S. Verstovsek, Genetic Analysis of Transforming Events That Convert Chronic Myeloproliferative Neoplasms to Leukemias. Cancer Research 70, 447–452 (2010).

10. F. Delhommeau, S. Dupont, V. D. Valle, C. James, S. Trannoy, A. Massé, O. Kosmider, J.-P. Le Couedic, F. Robert, A. Alberdi, Y. Lécluse, I. Plo, F. J. Dreyfus, C. Marzac, N. Casadevall, C. Lacombe, S. P. Romana, P. Dessen, J. Soulier, F. Viguié, M. Fontenay, W. Vainchenker, O. A. Bernard, Mutation in TET2 in Myeloid Cancers. N Engl J Med 360, 2289–2301 (2009).

11. A. Theocharides, M. Boissinot, F. Girodon, R. Garand, S.-S. Teo, E. Lippert, P. Talmant, Tichelli, S. Hermouet, R. C. Skoda, Leukemic blasts in transformed JAK2-V617F–positive myeloproliferative disorders are frequently negative for the JAK2-V617F mutation. Blood 110, 375–379 (2007).

12. P. A. Beer, F. Delhommeau, J.-P. LeCouédic, M. A. Dawson, E. Chen, D. Bareford, R. Kušec, M. F. McMullin, C. N. Harrison, A. M. Vannucchi, W. Vainchenker, A. R. Green, Two routes to leukemic transformation after a JAK2 mutation–positive myeloproliferative neoplasm. Blood 115, 2891–2900 (2010).

13. D. Rossi, C. Deambrogi, D. Capello, M. Cerri, M. Lunghi, G. Parvis, G. Saglio, G. Gaidano, D. Cilloni, JAK2 V ^617^ F mutation in leukaemic transformation of philadelphia-negative chronic myeloproliferative disorders. Br J Haematol 135, 267–268 (2006).

14. H. Celik, E. Krug, C. R. Zhang, W. Han, N. Issa, W. K. Koh, H. Bjeije, O. Kukhar, M. Allen, T. Li, D. A. C. Fisher, J. S. Fowles, T. N. Wong, M. C. Stubbs, H. K. Koblish, S. T. Oh, G. A. Challen, A Humanized Animal Model Predicts Clonal Evolution and Therapeutic Vulnerabilities in Myeloproliferative Neoplasms. Cancer Discovery 11, 3126–3141 (2021).

15. R. Rampal, J. Ahn, O. Abdel-Wahab, M. Nahas, K. Wang, D. Lipson, G. A. Otto, R. Yelensky, T. Hricik, A. S. McKenney, G. Chiosis, Y. R. Chung, S. Pandey, M. R. M. Van Den Brink, S. A. Armstrong, A. Dogan, A. Intlekofer, T. Manshouri, C. Y. Park, S. Verstovsek, F. Rapaport, P. J. Stephens, V. A. Miller, R. L. Levine, Genomic and functional analysis of leukemic transformation of myeloproliferative neoplasms. Proc. Natl. Acad. Sci. U.S.A. 111 (2014).

16. Y. Ushijima, S. Naruse, Y. Ishikawa, N. Kawashima, M. Sanada, M. Nakashima, J. H. Kim, S. Terakura, R. Kihara, K. Watamoto, T. Nishiyama, K. Kitamura, T. Matsushita, H. Kiyoi, Initiating-clone analysis in patients with acute myeloid leukemia secondary to essential thrombocythemia. Sci Rep 14, 15906 (2024).

17. K. Moran-Crusio, L. Reavie, A. Shih, O. Abdel-Wahab, D. Ndiaye-Lobry, C. Lobry, M. E. Figueroa, A. Vasanthakumar, J. Patel, X. Zhao, F. Perna, S. Pandey, J. Madzo, C. Song, Q. Dai, C. He, S. Ibrahim, M. Beran, J. Zavadil, S. D. Nimer, A. Melnick, L. A. Godley, I. Aifantis, R. L. Levine, Tet2 Loss Leads to Increased Hematopoietic Stem Cell Self-Renewal and Myeloid Transformation. Cancer Cell 20, 11–24 (2011).

18. F. Pan, T. S. Wingo, Z. Zhao, R. Gao, H. Makishima, G. Qu, L. Lin, M. Yu, J. R. Ortega, J. Wang, A. Nazha, L. Chen, B. Yao, C. Liu, S. Chen, O. Weeks, H. Ni, B. L. Phillips, S. Huang, J. Wang, C. He, G.-M. Li, T. Radivoyevitch, I. Aifantis, J. P. Maciejewski, F.-C. Yang, P. Jin, M. Xu, Tet2 loss leads to hypermutagenicity in haematopoietic stem/progenitor cells. Nat Commun 8, 15102 (2017).

19. L. Holmfeldt, C. G. Mullighan, The Role of TET2 in Hematologic Neoplasms. Cancer Cell 20, 1–2 (2011).

20. V. Santini, M. Stahl, D. A. Sallman, TP53 Mutations in Acute Leukemias and Myelodysplastic Syndromes: Insights and Treatment Updates. Am Soc Clin Oncol Educ Book 44, e432650 (2024).

21. N. G. Daver, A. Maiti, T. M. Kadia, P. Vyas, R. Majeti, A. H. Wei, G. Garcia-Manero, C. Craddock, D. A. Sallman, H. M. Kantarjian, TP53 -Mutated Myelodysplastic Syndrome and Acute Myeloid Leukemia: Biology, Current Therapy, and Future Directions. Cancer Discovery 12, 2516–2529 (2022).

22. P. Gou, D. Liu, S. Ganesan, E. Lauret, N. Maslah, V. Parietti, W. Zhang, V. Meignin, J.-J. Kiladjian, B. Cassinat, S. Giraudier, Genomic and functional impact of Trp53 inactivation in JAK2V617F myeloproliferative neoplasms. Blood Cancer J. 14, 1 (2024).

23. N. B. Leimkühler, H. F. E. Gleitz, L. Ronghui, I. A. M. Snoeren, S. N. R. Fuchs, J. S. Nagai, B. Banjanin, K. H. Lam, T. Vogl, C. Kuppe, U. S. A. Stalmann, G. Büsche, H. Kreipe, I. Gütgemann, P. Krebs, Y. Banz, P. Boor, E. W.-Y. Tai, T. H. Brümmendorf, S. Koschmieder, M. Crysandt, E. Bindels, R. Kramann, I. G. Costa, R. K. Schneider, Heterogeneous bone-marrow stromal progenitors drive myelofibrosis via a druggable alarmin axis. Cell Stem Cell 28, 637-652.e8 (2021).

24. L. F. Mendez Luque, A. L. Blackmon, G. Ramanathan, A. G. Fleischman, Key Role of Inflammation in Myeloproliferative Neoplasms: Instigator of Disease Initiation, Progression. and Symptoms. Curr Hematol Malig Rep 14, 145–153 (2019).

25. H. C. Hasselbalch, M. E. Bjørn, MPNs as Inflammatory Diseases: The Evidence, Consequences, and Perspectives. Mediators Inflamm 2015, 102476 (2015).

26. E. M. Soyfer, A. G. Fleischman, Myeloproliferative neoplasms – blurring the lines between cancer and chronic inflammatory disorder. Front. Oncol. 13, 1208089 (2023).

27. S. Koschmieder, N. Chatain, Role of inflammation in the biology of myeloproliferative neoplasms. Blood Reviews 42, 100711 (2020).

28. R. C. Zhang, D. Nix, M. Gregory, M. A. Ciorba, E. L. Ostrander, R. D. Newberry, D. H. Spencer, G. A. Challen, Inflammatory cytokines promote clonal hematopoiesis with specific mutations in ulcerative colitis patients. Experimental Hematology 80, 36-41.e3 (2019).

29. Wiley, T. M. Parsons, S. Burkart, A. L. Young, K. M. Erlandson, K. K. Tassiopoulos, K. Wu, C. Gurnett, R. M. Presti, K. L. Bolton, G. A. Challen, Effect of Clonal Hematopoiesis on Cardiovascular Disease in People Living with HIV. Experimental Hematology 114, 18–21 (2022).

30. B. Hormaechea-Agulla, K. A. Matatall, D. T. Le, B. Kain, X. Long, P. Kus, R. Jaksik, G. A. Challen, M. Kimmel, K. Y. King, Chronic infection drives Dnmt3a-loss-of-function clonal hematopoiesis via IFNγ signaling. Cell Stem Cell 28, 1428-1442.e6 (2021).

31. A. R. Zhang, E. L. Ostrander, O. Kukhar, C. Mallaney, J. Sun, E. Haussler, H. Celik, W. K. Koh, K. Y. King, P. Gontarz, G. A. Challen, Txnip Enhances Fitness of Dnmt3a -Mutant Hematopoietic Stem Cells via p21. Blood Cancer Discovery 3, 220–239 (2022).

32. J. M. SanMiguel, E. Eudy, M. A. Loberg, K. A. Young, J. J. Mistry, K. D. Mujica, L. S. Schwartz, T. M. Stearns, G. A. Challen, J. J. Trowbridge, Distinct Tumor Necrosis Factor Alpha Receptors Dictate Stem Cell Fitness versus Lineage Output in Dnmt3a -Mutant Clonal Hematopoiesis. Cancer Discovery 12, 2763–2773 (2022).

33. S. O. Abegunde, R. Buckstein, R. A. Wells, M. J. Rauh, An inflammatory environment containing TNFα favors Tet2-mutant clonal hematopoiesis. Experimental Hematology 59, 60–65 (2018).

34. C. Quin, E. N. DeJong, A. J. M. McNaughton, M. M. Buttigieg, S. Basrai, S. Abelson, M. J. Larché, M. J. Rauh, D. M. E. Bowdish, Chronic TNF in the aging microenvironment exacerbates Tet2 loss-of-function myeloid expansion. Blood Advances 8, 4169–4180 (2024).

35. Y. Wang, X. Zuo, Cytokines frequently implicated in myeloproliferative neoplasms. Cytokine X 1, 100005 (2019).

36. N. Chatain, S. Koschmieder, E. Jost, Role of Inflammatory Factors during Disease Pathogenesis and Stem Cell Transplantation in Myeloproliferative Neoplasms. Cancers 12, 2250 (2020).

37. H. Xiong, H. Zhang, J. Bai, Y. Li, L. Li, L. Zhang, Associations of the circulating levels of cytokines with the risk of myeloproliferative neoplasms: a bidirectional mendelian-randomization study. BMC Cancer 24, 531 (2024).

38. G. Trinchieri, Interleukin-12 and the regulation of innate resistance and adaptive immunity. Nat Rev Immunol 3, 133–146 (2003).

39. S. S. Cho, C. M. Bacon, C. Sudarshan, R. C. Rees, D. Finbloom, R. Pine, J. J. O’Shea, Activation of STAT4 by IL-12 and IFN-alpha: evidence for the involvement of ligand-induced tyrosine and serine phosphorylation. J Immunol 157, 4781–4789 (1996).

40. W. T. Watford, B. D. Hissong, J. H. Bream, Y. Kanno, L. Muul, J. J. O’Shea, Signaling by IL-12 and IL-23 and the immunoregulatory roles of STAT4. Immunological Reviews 202, 139–156 (2004).

41. W. E. Thierfelder, J. M. Van Deursen, K. Yamamoto, R. A. Tripp, S. R. Sarawar, R. T. Carson, M. Y. Sangster, D. A. A. Vignali, P. C. Doherty, G. C. Grosveld, J. N. Ihle, Requirement for Stat4 in interleukin-12-mediated responses of natural killer and T cells. Nature 382, 171–174 (1996).

42. M. J. Slade, R. Ghasemi, M. O’Laughlin, T. Burton, R. S. Fulton, H. J. Abel, E. J. Duncavage, T. J. Ley, M. A. Jacoby, D. H. Spencer, Persistent Molecular Disease in Adult Patients With AML Evaluated With Whole-Exome and Targeted Error-Corrected DNA Sequencing. JCO Precis Oncol 7, e2200559 (2023).

43. A. L. Young, H. C. Davis, M. J. Cox, T. M. Parsons, S. C. Burkart, D. E. Bender, L. Sun, S. T. Oh, G. A. Challen, Spatial Mapping of Hematopoietic Clones in Human Bone Marrow. Blood Cancer Discovery 5, 153–163 (2024).

44. T. M. Parsons, A. Krishnan, W. M. B. Dunuwille, A. L. Young, J. Arand, W. Han, G. A. Challen, Engineering a humanized animal model of polycythemia vera with minimal JAK2V617Fmutant allelic burden. haematol, doi: 10.3324/haematol.2023.283858 (2023).

45. K. Clement, H. Rees, M. C. Canver, J. M. Gehrke, R. Farouni, J. Y. Hsu, M. A. Cole, D. R. Liu, J. K. Joung, D. E. Bauer, L. Pinello, CRISPResso2 provides accurate and rapid genome editing sequence analysis. Nat Biotechnol 37, 224–226 (2019).

46. A. Mullally, S. W. Lane, B. Ball, C. Megerdichian, R. Okabe, F. Al-Shahrour, M. Paktinat, J. E. Haydu, E. Housman, A. M. Lord, G. Wernig, M. G. Kharas, T. Mercher, J. L. Kutok, D. G. Gilliland, B. L. Ebert, Physiological Jak2V617F expression causes a lethal myeloproliferative neoplasm with differential effects on hematopoietic stem and progenitor cells. Cancer Cell 17, 584–596 (2010).

47. S. Xiong, D. Chachad, Y. Zhang, J. Gencel-Augusto, M. Sirito, V. Pant, P. Yang, C. Sun, G. Chau, Y. Qi, X. Su, E. M. Whitley, A. K. El-Naggar, G. Lozano, Differential Gain-of-Function Activity of Three p53 Hotspot Mutants In Vivo. Cancer Res 82, 1926–1936 (2022).

48. Z. Xie, A. Bailey, M. V. Kuleshov, D. J. B. Clarke, J. E. Evangelista, S. L. Jenkins, A. Lachmann, M. L. Wojciechowicz, E. Kropiwnicki, K. M. Jagodnik, M. Jeon, A. Ma’ayan, Gene Set Knowledge Discovery with Enrichr. Current Protocols 1, e90 (2021).

49. J. B. Clarke, M. Jeon, D. J. Stein, N. Moiseyev, E. Kropiwnicki, C. Dai, Z. Xie, M. L. Wojciechowicz, S. Litz, J. Hom, J. E. Evangelista, L. Goldman, S. Zhang, C. Yoon, T. Ahamed, S. Bhuiyan, M. Cheng, J. Karam, K. M. Jagodnik, I. Shu, A. Lachmann, S. Ayling, S. L. Jenkins, A. Ma’ayan, Appyters: Turning Jupyter Notebooks into data-driven web apps. Patterns 2, 100213 (2021).

50. B. Ruffell, D. Chang-Strachan, V. Chan, A. Rosenbusch, C. M. T. Ho, N. Pryer, D. Daniel, E. S. Hwang, H. S. Rugo, L. M. Coussens, Macrophage IL-10 Blocks CD8+ T Cell-Dependent Responses to Chemotherapy by Suppressing IL-12 Expression in Intratumoral Dendritic Cells. Cancer Cell 26, 623–637 (2014).

