## Supplemental Materials for "*JAK2*^V617F^ Myeloproliferative Neoplasms Support Parallel Evolution of Independent Leukemic Clones"

**The PDF file includes:**

Materials and Methods
Figs. S1 to S8
Tables S1 to S2
References 42-50

**Other Supplementary Materials for this manuscript include the following:**

Data S1

### Materials and Methods

#### *Human Samples*

All cord blood specimens were anonymized and no member of the study team had access to information that allowed the specimens to be linked to identifiable individuals. These studies were thus deemed "nonhuman studies" by the Washington University Human Research Protection Office. De-identified cord blood specimens were collected as part of a study approved by the Human Research Protection Office and the Institutional Review Board at Washington University School of Medicine (IRB# 202104011) after patients provided informed consent in accordance with the Declaration of Helsinki. De-identified bone marrow sample specimens were collected as healthy donor controls (IRB# 201103258). MPN patient samples were obtained according to a protocol approved by the Washington University Human Studies Committee (WU no. 01-1014). All patients previously provided consent to have samples banked and were not newly recruited for this study. Mutation information for patient samples was originally obtained from clinical testing with a targeted, error-corrected sequencing assay that covers 56 genes recurrently mutated in myeloid cancers. The assay has a documented limit of detection of 2% variant allele fraction (VAF) for new variants identified at initial diagnosis and 0.1% VAF for previously identified variants for molecular disease monitoring(*42*). Mononuclear cells using SepMate-50 (StemCell Technologies #85450) mediated ficoll gradient extraction according to standard procedures. CD34+ cells were isolated using magnetic enrichment (Miltenyi Biotec #130-100-453) and cultured overnight in SFEMII media (StemCell Technologies #09605) supplemented with 50 U/mL penicillin-streptomycin (Fisher Scientific #MT30002CI), 50 ng/mL human stem cell factor (SCF; Miltenyi Biotec #130–096–695), 50 ng/mL human thrombopoietin (TPO; Miltenyi Biotec #130–094–013), and 50 ng/mL human FLT3L (Miltenyi Biotec #130–096–479). Enrichment efficiency was confirmed by flow cytometry.

#### *Single Cell DNA Sequencing and Analysis*

Single-cell sequencing was performed using the Mission Bio Tapestri platform using methods similar to previous single-cell studies(*43*).Cryovials of patient bone marrow or peripheral blood were thawed, resuspended in Hanks Balanced Salt Solution (HBBS, Corning #21021CV) and quantified by cellometer (Nexelom Bioscience). Cells were stained with 7AAD and live cells were sorted using a MoFlo cell sorter (Dako). One million live cells per sample were utilized for single-cell analysis on the Tapestri platform. Live cells were encapsulated into microfluidic droplets using the Tapestri instrument, lysed, and barcoded for library amplification using the Mission Bio myeloid panel (Table S2). This panel covers 45 genes that are recurrently mutated in myeloid malignancies. After library amplification the droplets were broken, and the libraries were isolated by Ampure XP bead cleanup (Beckman Coulter). Each purified library underwent PCR amplification using sequencing primers containing library-specific indexes followed by Ampure XP bead cleanup. Library quality was assessed using the TapeStation (Agilent Technologies) and Qubit (Thermo Fisher Scientific). Libraries were sequenced on the Illumina NovaSeq 6000 platform using the 300-cycle kit targeting 173 million read pairs for the DNA myeloid panel to ensure adequate coverage across the panel for each cell captured. FASTQ files containing the sequenced read information were analyzed using the cloud-based Tapestri Pipeline without modification. The pipeline trimmed the adapter sequences, mapped reads to the human reference genome (hg19) using BWA, and assigned each read to a unique cell. GATK v4/Haplotypecaller was used to call genotypes for each cell. Somatic mutations were retained if they were observed in >1% of cells and were not likely to arise due to allelic dropout. Sequencing results were further normalized, clustered, and annotated using the Mission Bio Mosaic version 2.4. Figures were generated using the Mission Bio Mosaic v2.4 pipeline.

*Droplet Digital PCR (ddPCR)*

*JAK2*$^{V617F}$ variant validation by ddPCR was performed on the QX200 platform (Bio-Rad) following similar methods as previously described(*44*). Briefly, 20-50 ng of DNA was utilized per individual reaction. Droplets were generated using the QX200 droplet generator (Bio-Rad) following standard manufacturer protocols. PCR amplification of droplets was performed in a deep-well thermocycler (Bio-Rad) followed by analysis on the QX200 droplet reader (Bio-Rad).

*CRISPR and VAF Determination*

Single guide RNAs (gRNA) targeting *TET2*, *TP53*, and *AAVS1* were designed using the UCSC Genome Browser. Multiple gRNAs were tested, the sequences for gRNAs (Synthego) used for final experimentation are as follows:

g*TET2*: GAAGCTACTGTGTTTGGTGC
g*TP53*: TGATGGTGAGGATGGGCCTC
g*AAVS1*: GGGGCCACTAGGGACAGGAT

Single stranded oligo donor nucleotides (ssODN) for *TET2*$^{R1216*}$, *TP53*$^{R248Q}$, and *AAVS1*$^{Silent}$ were designed using the UCSC Genome Browser and were purchased from IDT and are as follows:

*TET2*$^{R1216*}$:
GGTTGGGGTGGGGGGTGTTTGGGATGGAATGGTGATCCACGCAGGTGGTTCGCAGA
AGCAGCAGTGAAGAGAAGCTACTGTGTTTGGTGCGAGAGTGAGCTGGCCACACCTG
TGAGGCTGCAGTGATTGTGATTCTCATCCTGGTGTGGGAAGGAATCCC

*TP53*$^{R248Q}$:
GCCAGTGTGCAGGGTGGCAAGTGGCTCCTGACCTGGAGTCTTCCAGTGTGATGATGG
TGAGGATGGGCCTCTGGTTCATGCCGCCCATGCAGGAACTGTTACACATGTAGTTGT
AGTGGATGGTGGTACAGTCAGAGCCAACCTAGGAGATAACACAGGC

*AAVS1*$^{Silent}$:
AATGTGGCTCTGGTTCTGGGTACTTTTATCTGTCCCCTCCACCCCACAGTGGGGCCAC
TAGGGACAGGTTTGGTGACAGAAAAGCCCCATCCTTAGGCCTCCTCCTTCCTAGTCT
CCTGATATTGGGTCTAACCCCCACCTCC

Nucleofection was performed using a Neon Transfection System (Invitrogen #MPK5000) with the following parameters: Voltage: 1600, Width: 10, Pulses: 3. 48 hours post-nucleofection, approximately 100,000 cells were set aside to quantify CRISPR/Cas9 targeting efficiency using PCR amplicon-based deep sequencing. Libraries were sequenced with the Illumina MiSeq platform and output was analyzed using the CRISPResso2 web-based software(*45*). The following primer pairs were used to generate amplicons:

TET2 FWD: TGCAAGTGACCCTTGTTTTG
TET2 REV: ATTTCCTCAGCGTCTCGGTA

TP53 FWD: TGGAAGAAATCGGTAAGAGGTG
TP53 REV: TGGCTCTGACTGTACCACCA
AAVS1FWD: ACAGGAGGTGGGGGTTAGAC
AAVS1REV: CCCCTATGTCCACTTCA

*Ex Vivo Competition Assays*

CD34+ cells were immunomagnetically isolated (Miltenyi Biotec #130-046-702) using the AutoMACS Neo system from PV or MF patient specimens and healthy donor BM and used as "host" cells. In parallel, CD34+ cells were isolated from umbilical cord blood obtained from healthy donors and used as "test" cells. Test cells were nucleofected with gRNA and ssODN templates targeting TET2, TP53, or AAVS1 as above. Host and test cells were combined at a ratio of 100,000 host cells to 20,000 test cells per well in a round-bottom 96-well plate and maintained in SFEMII media (StemCell Technologies #09605) supplemented with 50 U/mL penicillin-streptomycin (Fisher Scientific #MT30002CI), 50 ng/mL human stem cell factor (SCF; Miltenyi Biotec #130–096–695), 50 ng/mL human thrombopoietin (TPO; Miltenyi Biotec #130–094–013), and 50 ng/mL human FLT3L (Miltenyi Biotec #130–096–479) under normoxic conditions. Co-cultures were harvested on days 6 and 12 post-plating. At each time point, cell pellets were collected for genomic DNA extraction using PureLink gDNA column-based purification kit (Invitrogen #K182002). Targeted loci were amplified by PCR and amplicons were sequenced and analyzed as above enabling assessment of relative clonal expansion of gene-edited test cells within the mixed culture.

*Mice and Transplantation*

The Institutional Animal Care and Use Committee at Washington University School of Medicine approved all animal procedures. Mice were housed in specific pathogen-free conditions at Washington University School of Medicine on a 12:12-hour light:dark cycle in temperature- and humidity-controlled rooms. Donor mice were typically 10-12-weeks old for experimentation. Both male and female mice were used. Mice used in murine competition experiments were all C57Bl/6 background. *Jak2*$^{V617F/+}$ (*46*) and *Tet2*$^{fl/+}$ (*17*) were crossed to Vav-Cre mice (The Jackson Laboratory #004682). *Tp53*$^{R172H/+}$ (*47*) and *Tnfrsf1b*$^{tm1Imx}$ *Tnfrsf1a*$^{tm1Imx}$ (*Tnfr*$^{KO}$; The Jackson Laboratory #003243) mice were germline knock-in and knock-out mutants respectively. Vav-Cre;*Tet2*$^{fl/+}$ mice were crossed to *Tnfr*$^{KO}$ to generate *Tet2*$^{\Delta/+}$*Tnfr*$^{KO}$ mice. Recipient mice (C57Bl/6 CD45.1, The Jackson Laboratory #002014) were approximately 8-weeks old and transplanted by retro-orbital injection after a split dose (4-hours apart) of 10.5 Gy irradiation. For two-way chimeras, $2.5x10^6$ whole bone marrow (WBM) cells from *Jak2*$^{V617F}$ or WT control (host, CD45.2) were transplanted along with $5x10^5$ WBM cells from either *Tet2*$^{\Delta/+-}$, *Tp53*$^{R172H/+}$, or WT control (Test, CD45.1/2). For the *Jak2*$^{V617F}$ titration experiment, a range of WBM ($2.5x10^6$, $1.25x10^6$, $6.25x10^5$, $3.12x10^5$, $1.56x10^5$) sourced from *Jak2*$^{V617F}$ (host, CD45.2) was transplanted with a range of WBM ($1.25x10^6$, $1.88x10^6$, $2.19x10^6$, $2.34x10^6$, $2.50x10^6$) sourced from WT (support, CD45.1), and $5.0x10^5$ WBM cells sourced from *Tet2*$^{\Delta/+}$ (test, CD45.1/2). For three-way chimeras after the titration experiment, $1.5x10^6$ WBM cells from *Jak2*$^{V617F}$ (host, CD45.2) was transplanted with $1.05x10^6$ WBM cells from WT (support, CD45.1) and $4.5x10^5$ WBM cells from *Tet2*$^{\Delta/+}$ (test, CD45.1/2). For TNF-receptor knock-out studies, chimeras were established with the same WBM cell numbers as in the three-way chimera experiments with *TNFr*$^{KO}$ and *Tet2*$^{\Delta/+}$*Tnfr*$^{KO}$ as test (CD45.2), *Jak2*$^{V617F}$ or WT as host (CD45.1/2), and WT as support (CD45.1).

*PDX Transplantation*

For patient derived xenograft (PDX) experiments, NOD-scid- *Il2rg*-null-3/GM/SF (NSGS; The Jackson Laboratory #013062) were used as recipients. $2.0 \times 10^4$ cord-blood derived $CD34^+$ cells from *AAVS1*, *TET2* and *TP53* nucleofected populations were transplanted with $1.0 \times 10^5$ $CD34^+$ cells derived from healthy donor (HD) human BM or PV or MF patient samples into sublethally irradiated (250 rads) 6–8-week-old NSGS mice via intra-tibial injections in a volume of 30 µL with 29-gauge U-100 insulin syringes (Covetrus #074076). For intra-tibial injections, mice were anesthetized with an intra-muscular injection of Ketamine/Xylazine mixture (2 mg/mouse; KetaVed, Vedco). The needle was inserted approximately 0.8 cm deep into the tibia and cells were gradually released into BM while the needle was gently removed.

*Cell Purification and Flow Cytometry of Cell Surface Markers*

BM cells were isolated from iliac crests, femurs, and tibias of mice. Cells were stained in Hanks Balanced Salt Solution (HBBS, Corning #21021CV) containing 100 U/mL penicillin/streptomycin (Fisher Scientific #MT30002CI), 10 umol/L HEPES (Life Technologies #15630080) and 2% Serum Plus II (Sigma #14009C) at a density of $1.0 \times 10^8$/mL. Staining was performed for >30 minutes at 4°C with desired antibodies. For cell sorting, BM was enriched with anti-mouse CD117-conjugated microbeads (Miltenyi Biotec #130-0910224) using the AutoMacs Neo Separator (Miltenyi Biotec), then stained with appropriate antibody cocktails. For HSC analysis and HSPC sorting from BM, the following antibodies were used: CD45.1-BV785 (clone A20; BioLegend #110743), CD45.2-BV605 (clone 104; BioLegend #109841), B220-APCcy7 (clone RA3-6B2; BioLegend #103224), Gr-1-APCcy7 (clone RB6-8C5; BioLegend #108424), Mac-1-APCcy7 (clone M1/70; BioLegend #101226), CD3e-APCcy7 (clone 145-2C11; BioLegend #100330), Ter119-APCcy7 (clone TER-119; BioLegend #116223), CD48-PECy7 (clone HM48-1; BioLegend #103424), CD150-PE (clone TC15-12F12.2; BioLegend #115904), c-Kit-BV421 (clone 2B8; BioLegend #105828), Sca1-APC (clone E13-161.7; BioLegend #122512).

For mouse peripheral blood analysis, blood samples were obtained by venipuncture. Red blood cells were lysed then samples were stained with the following antibodies: CD45.1-PE (clone A20; BioLegend #110708), CD45.2-BV421 (clone 104; BioLegend #109831), B220-PECy7 (clone RA3-6B2; BioLegend #103222), B220-APCCy7 (clone RA3-6B2; BioLegend #103224), Gr-1-PECy7 (clone RB6-8C5; BioLegend #108416), CD11b-PECy7 (clone M1/70; BioLegend #101216), CD3e-APCCy7 (clone 145-2C11; BioLegend #100330)

For PDX systems, the following antibodies were used for PB and BM sorting and analysis: anti-mouse CD45-BV605 (clone 30-F11; BioLegend #103139), anti-human CD45-APC (clone 2D1; BioLegend #368512), anti-human CD3-PEcy7 (clone HIT3a; BioLegend #300316), anti-human CD19-FITC (clone 4G7; BioLegend #392508), anti-human CD33-BV421 (clone WM53; BD #565949), anti-human CD34-PE (clone 561; BioLegend #343606), anti-human CD90-PECy7 (clone 5E10; BioLegend #328124), anti-human CD45RA-BV421 (clone HI100; BioLegend #304130), anti-human CD38-FITC (clone HB7; Invitrogen #11-0388-42), anti-human Lineage cocktail-BV510 (BioLegend #348807), and anti-human CD45-biotin (clone HI30; BioLegend #304004).

Dead cells were excluded from all analyses and downstream applications with 7AAD (BioLegend #420404, 1:100 dilution). Cell sorting was performed using MoFlo (Beckman Coulter). Flow cytometric analysis was performed using Northern Lights (CyTek). Acquired flow cytometry data were analyzed with FlowJo software.

*RNA Sequencing*

7,000 donor-derived HSPCs (Lineage$^{-}$ Sca-1$^{+}$ c-Kit$^{+}$ = "LSK") from primary transplants were isolated using MoFlo (Beckman Coulter) from two pooled biological replicates from two independent cohorts (four total replicates per genotype per condition). Total RNA was extracted using NucleoSpin RNA Plus XS kit (Takara Bio #740990.50) and RNA Integrity was determined using Agilent Bioanalyzer. RNA with a RIN >8.0 was used to prepare for cDNA library with the SMARTer Ultra Low RNA kit for Illumina Sequencing (Takara-Clontech) per manufacturer's protocol. Fragments were sequenced on an Illumina NovaSeq X Plus using paired end reads extending 150 bases.

*RNA Sequencing Analysis*

Basecalls and demultiplexing were performed with Illumina's DRAGEN and BCLconvert version 4.2.4 software with a maximum of one mismatch in the indexing read. RNA-seq reads were then aligned to the Ensembl release 101 primary assembly with STAR version 2.7.9a. Gene counts were derived from the number of uniquely aligned unambiguous reads by Subread:featureCount version 2.0.3. Sequencing performance was assessed for the total number of aligned reads, total number of uniquely aligned reads and features detected. Gene counts were imported into the R studio and used EdgeR and TMM packages to analyze normalization size factors to adjust for sample differences in library size. The matrix counts and TMM size factors were imported using Limma package. The mean-variance relationship of every gene and sample was then calculated and the count matrix was converted to log2 counts-per-million using Limma's voom. Differential gene expressions (DEG) were analyzed using Limma with Benjamini-Hochberg false discovery rate. DEGs were defined by filtering for unadjusted $p$-value=0.05 and logFC=0.58 (equivalent to 1.5-fold change). Over-representation analysis of cancer-related pathways was performed using the NCI-Nature cancer pathways gene set library. Data visualization was performed using Enrichr and Appyters web-based software application package of meta Jupyter Notebook executions (volcano plot of ORA and UMAP of ORA) and MaGIC Volcano Plot Tool Software for volcano plot of DEGs(*48*, *49*).

*pSTAT4 Intracellular Flow Cytometry*

BM cells were stained for cell surface markers using the above makers in Cell Staining Buffer (BioLegend #420201) supplemented with dissolved Pierce Phosphatase Inhibitor tablets (Thermo Scientific #A32957). Following surface marker staining, whole BM was lysed and fixed with RBC Lysis/Fixation Solution (BioLegend #422401) at 37C for 15 minutes and subsequent permeabilization with True Phos Perm Buffer (BioLegend #425401) at -20ºC for >60 minutes. After washing, the cells were resuspended at a concentration of 10x10$^{6}$ cells per mL of buffer. 100uL of this cell solution was stained with anti-mouse pSTAT4-PE (clone A19016A; BioLegend #941206) at room temperature for 30 minutes in the dark. Flow cytometric analysis was performed using Northern Lights (CyTek). Acquired flow cytometry data were analyzed with FlowJo Software.

*Serum Cytokine Analysis*

Peripheral blood samples were obtained by venipuncture and collected into serum separator tubes (BD Microtainer SST #365967). Samples were allowed to clot at room temperature for 30 minutes followed by centrifugation at 1,500g for 10 minutes at 4°C. Serum was stored at -80°C until analysis. Frozen serum aliquots were shipped to Eve Technologies Corporation (Calgary, AB, Canada) for multiplex cytokine analysis (Mouse Cytokine/ Chemokine 32-Plex Discovery Assay Array or Human Cytokine/Chemokine 48-Plex Discovery Assay Array). Assays were performed according to the manufacturer's protocols using a bead-based immunoassay (Luminex xMAP technology). Data were acquired via a Luminex 200 system, and analyte concentrations were calculated from standard curves generated with referenced cytokine standards.

*in Vitro Cytokine Assays*

$CD34^+$ cells were isolated from umbilical cord blood obtained from healthy donors and used as "test" cells. Test cells were nucleofected with gRNA and ssODN templates targeting *TET2* as above. 100,000 nucleofected cells were plated per well in a round-bottom 96-well plate and maintained in SFEMII media (StemCell Technologies #09605) supplemented with 50 U/mL penicillin-streptomycin (Fisher Scientific #MT30002CI), 100 ng/mL human stem cell factor (SCF; Miltenyi Biotec #130–096–695), 100 ng/mL human thrombopoietin (TPO; Miltenyi Biotec #130–094–013), and 100 ng/mL human FLT3L (Miltenyi Biotec #130–096–479) under normoxic conditions. Recombinant human IL-27 (PeproTech #200-38), IL1b (PeproTech #200-011B), IL-12 (PeproTech #200-12H), IL-1RA (PeproTech #200-01RA), or TNFα (PeproTech #300-01A) was added at concentrations of 0, 1, 25, 50, and 100 ng/mL each into separate wells containing CD34+ cells every 48 hours. Co-cultures were harvested on day 6 post-plating and cell pellets were collected for genomic DNA extraction using PureLink gDNA column-based purification kit (Invitrogen #K182002). Targeted loci were amplified by PCR and amplicons were sequenced and analyzed as above enabling assessment of relative clonal expansion of gene-edited test cells tested at each cytokine and each concentration.

*in vivo Cytokine Neutralization*

Mouse neutralizing InVivoMAb antibodies against interleukin-12 (Bio X Cell #BE0052) and TNFα (Bio X Cell #BE0058) and the human neutralizing InVivoSIM antibody against interleukin-12 (Bio X Cell #SIM0020) were administered *in vivo*. Mice were randomized into treatment groups based on test cell peripheral blood engraftment. Neutralizing antibodies were administered intraperitoneally at 500ug per injection twice per week (cumulative weekly dose of 1mg) for a total treatment duration of 5-weeks during each transplant period as adopted and modified from prior studies(*50* ). Control animals received isotype-matched antibodies following the same dosing schedule. All injections were performed under aseptic technique with mice closely monitored for signs of distress throughout the study period.

*Quantitative Real-time PCR*

HSPCs ($Lineage^-$ $Sca\text{-}1^+$ $c\text{-}Kit^+$ = "LSK") were isolated using MoFlo (Beckman Coulter) from three pooled biological replicates. Total RNA was extracted using NucleoSpin RNA Plus XS kit (Takara Bio #740990.50). cDNA was synthesized using SuperScript VILO cDNA kit (ThermoFisher Scientific #1174050) and diluted 1:10 for real-time qPCR assays. Diluted cDNA samples were used for qPCR with Taq Man Master Mix (ThermoFisher Scientific #4304437), probes, and 18S (internal control). qPCR was performed using a Step One Plus Real-Time qPCR machine. The following probes (ThermoFisher Scientific #4331182) were used for the assay: *Hoxa10* (Mm00433966_m1), *Hlf* (Mm00723157_m1), *Fgd5* (Mm00554954_m1), *Mecom* (Mm00491303_m1), *Mki67* (Mm01278617_m1), *S100a8* (Mm00496696_g1), and *S100a9* (Mm00656925_m1).

*Statistical Analysis*

One-way ANOVA with Tukey correction for multiple comparisons (multiple groups) or unpaired two-tailed t-test (two groups) were used for statistical comparison where appropriate. Significance is indicated using the following convention: *$p \leq 0.05$, **$p \leq 0.01$, ***$p \leq 0.001$, ****$p \leq 0.0001$. All graphs represent mean ± s.e.m. unless otherwise indicated.

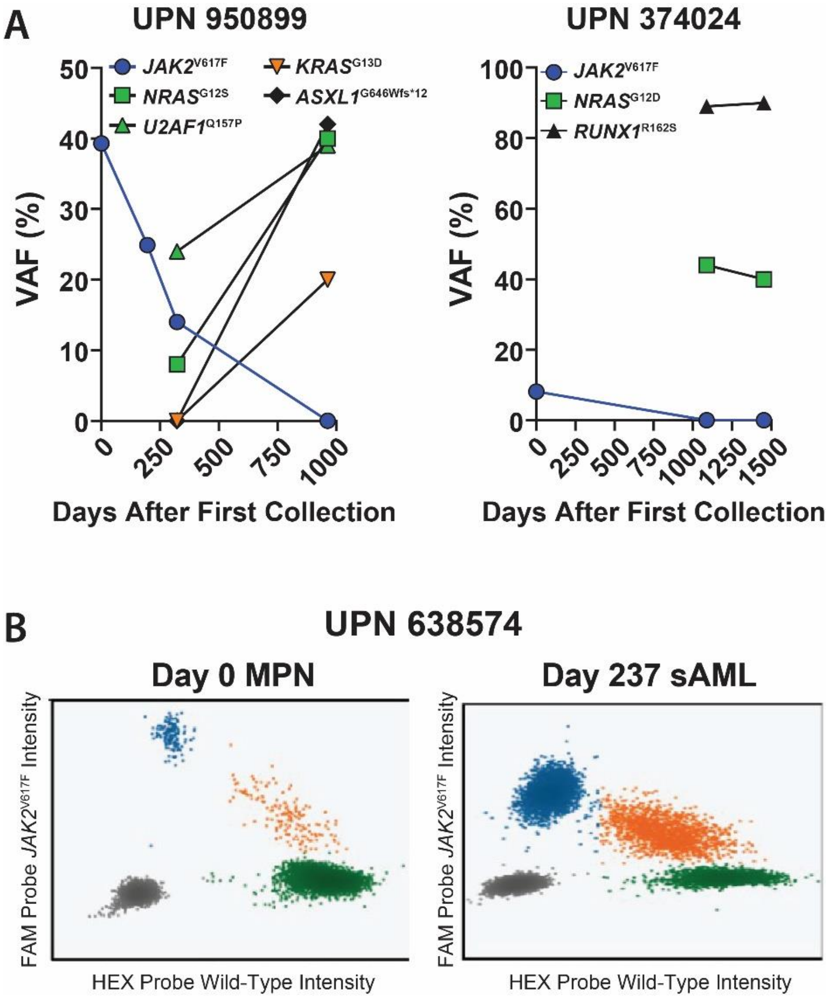


**Fig. S1.**

**(A)** Variant allele frequencies (VAF) of mutations identified from clinical sequencing for indicated patients. **(B)** ddPCR quantification of *JAK2*$^{V617F}$ mutant burden at MPN and sAML collection timepoints for UPN:638574.

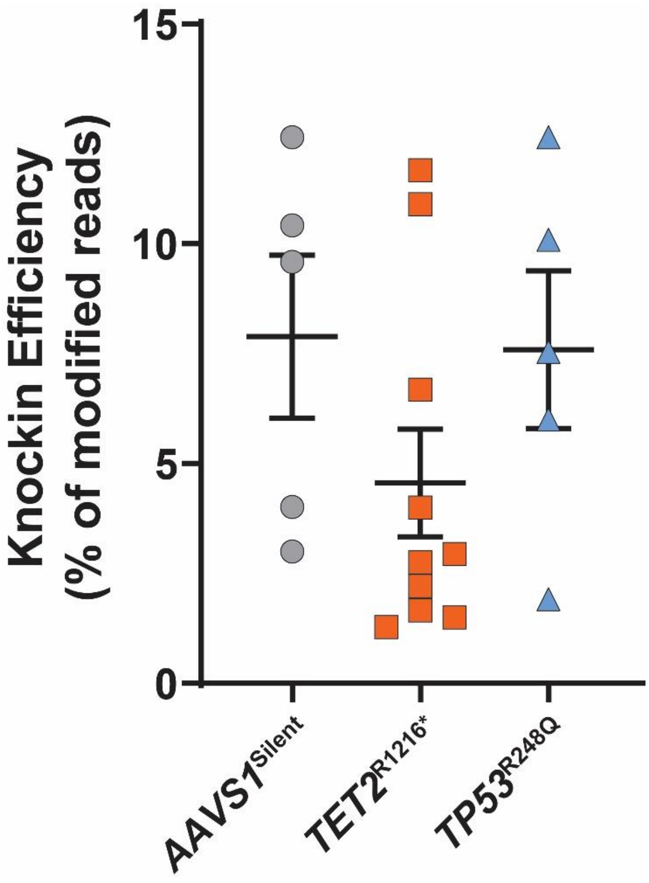


**Fig. S2.**
Starting CRISPR/Cas9 knock-in efficiencies of indicated mutations in cord blood $CD34^{+}$ cells determined by next-generation sequencing.

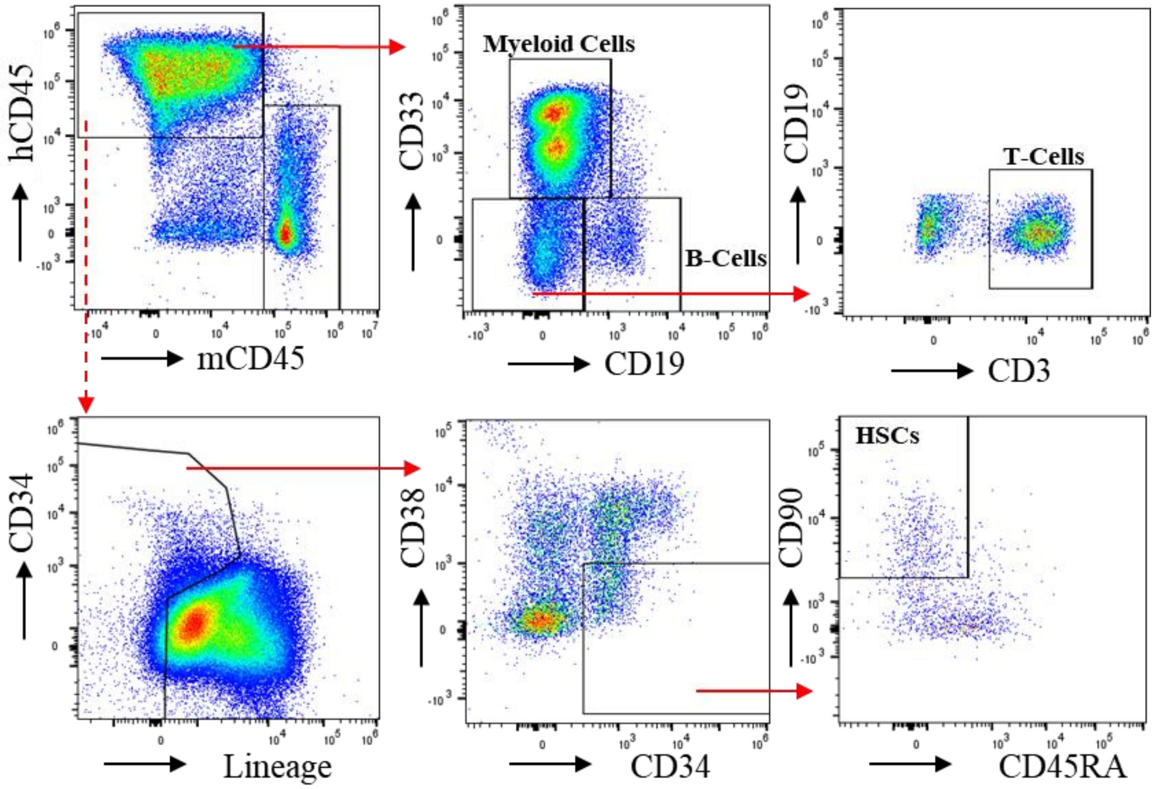


**Fig. S3.**
Representative flow cytometry gating scheme of live singlets to identify human cell engraftment, blood cell lineage and HSC populations in PDX experiments.

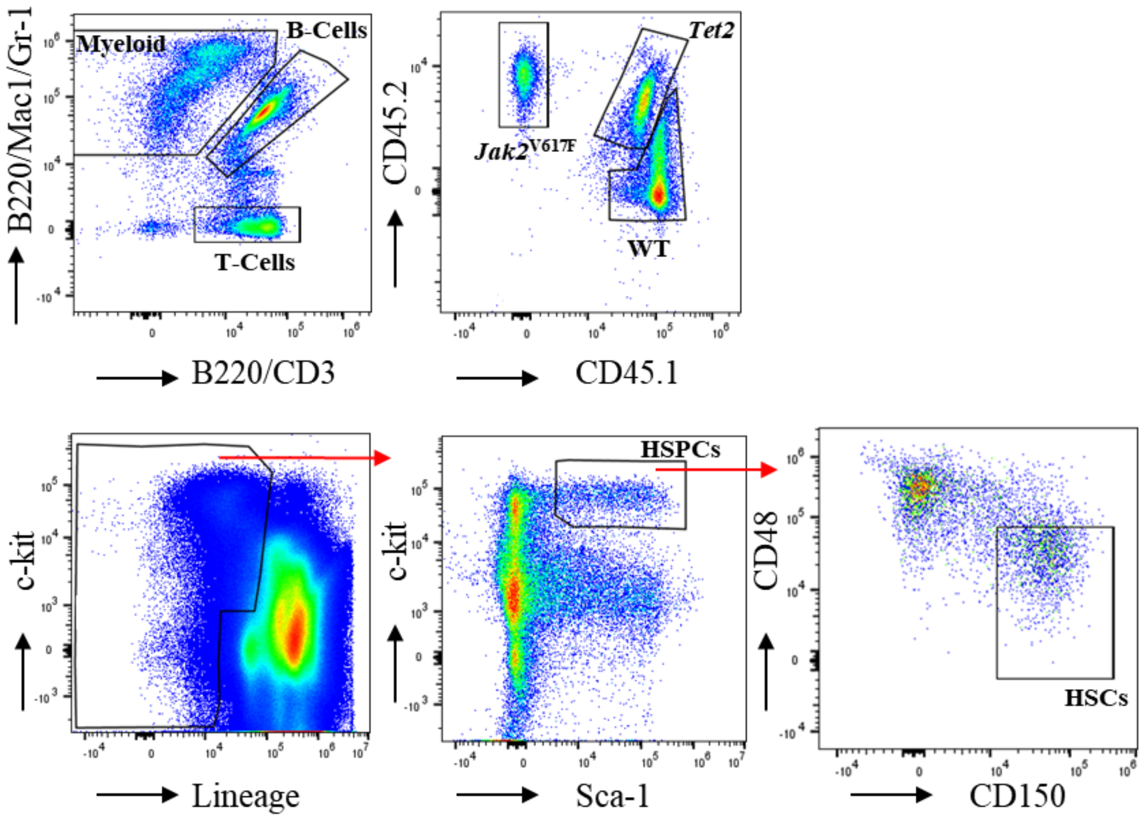


**Fig. S4.**
Representative flow cytometry gating scheme of live singlets to identify (top row) mouse peripheral blood cell lineage distribution and donor-derived chimerism and (bottom row) discrimination of mouse HSPC and HSC populations in BM of recipient mice.

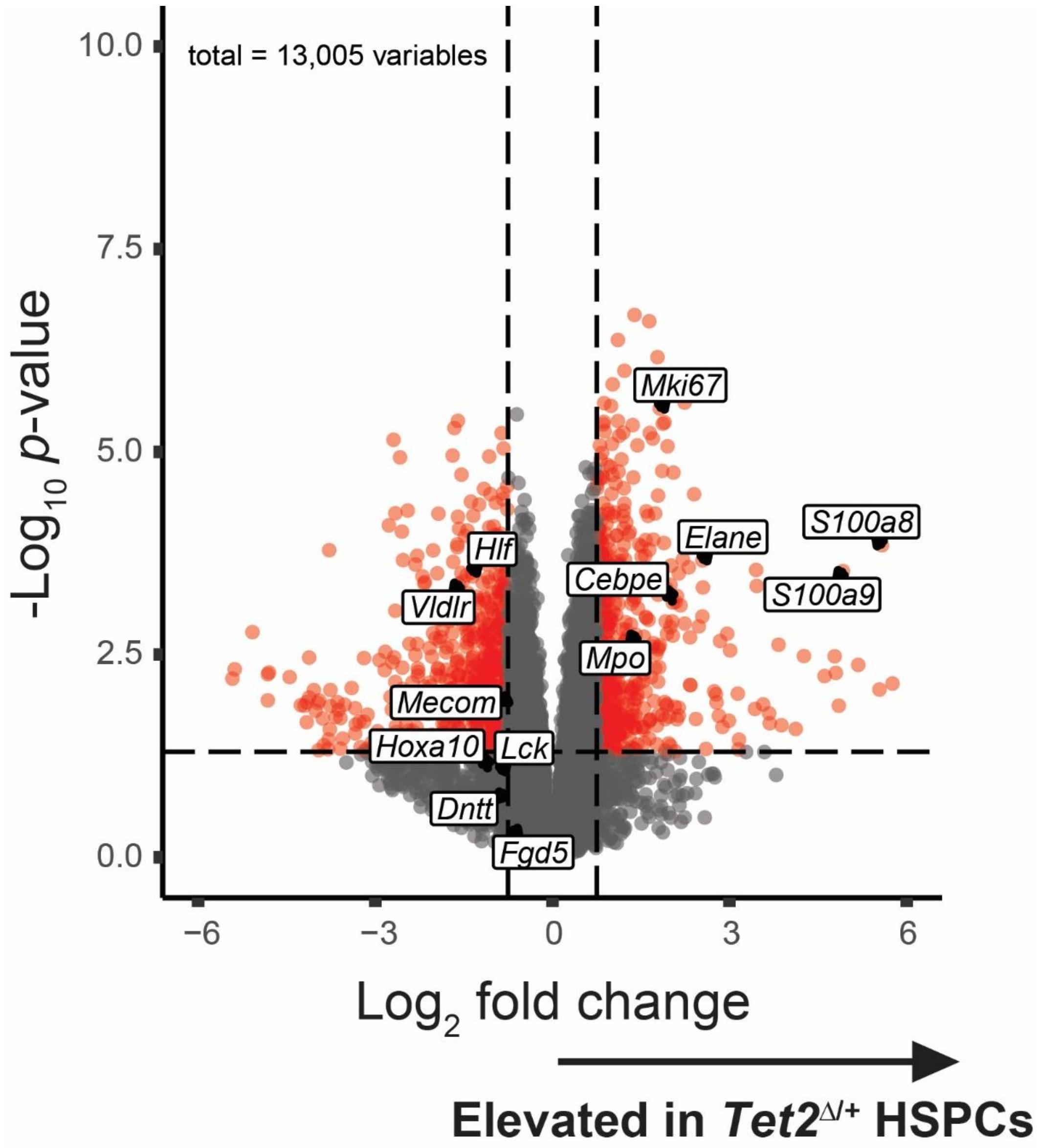


**Fig. S5.**

Volcano plot depicting differentially expressed genes between *Tet2*$^{\Delta/+}$ HSPCs from a *Jak2*$^{V617F}$ environment compared to WT HSPCs from a *Jak2*$^{V617F}$ environment.

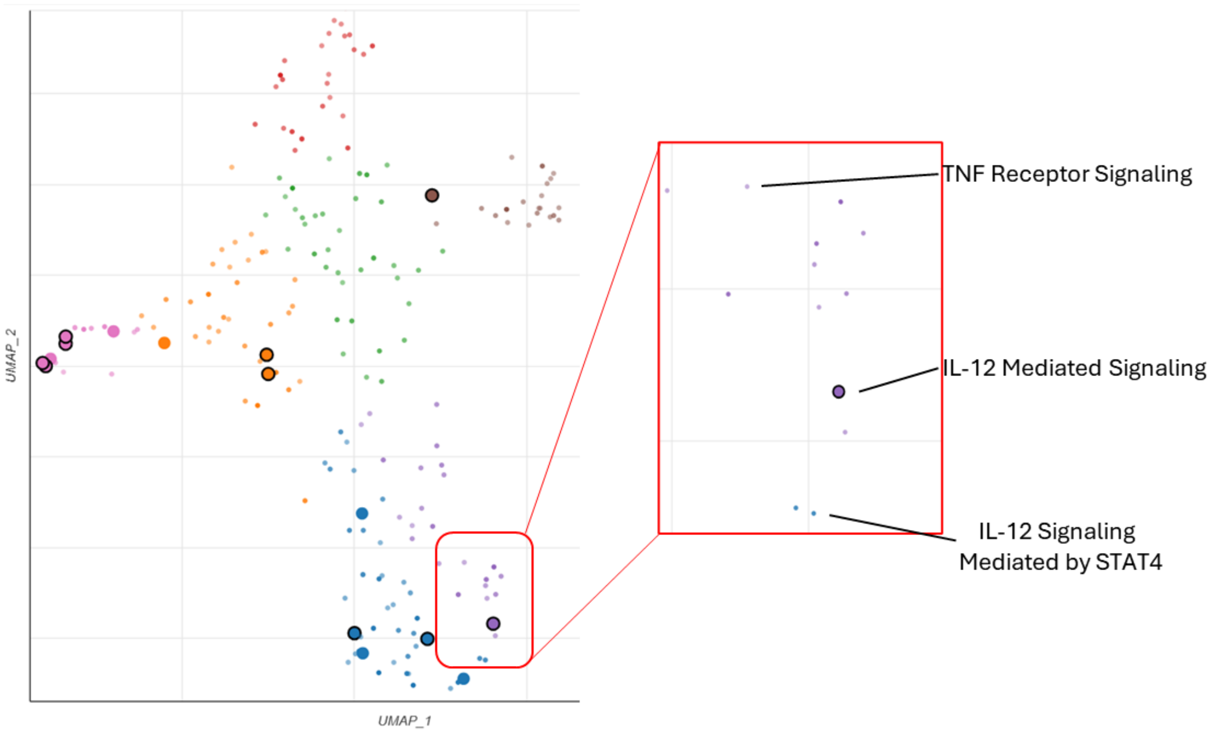


**Fig. S6.**

UMAP visualization of relationships between enriched cancer-related pathways determined by input list of differentially expressed genes between *Tet2*$^{\Delta/+}$ HSPCs cells from a *Jak2*$^{V617F}$ environment compared to WT HSPCs form a *Jak2*$^{V617F}$ environment. Terms with more similar gene sets are positioned closer together. The darker and larger the point, the more significantly enriched the term.

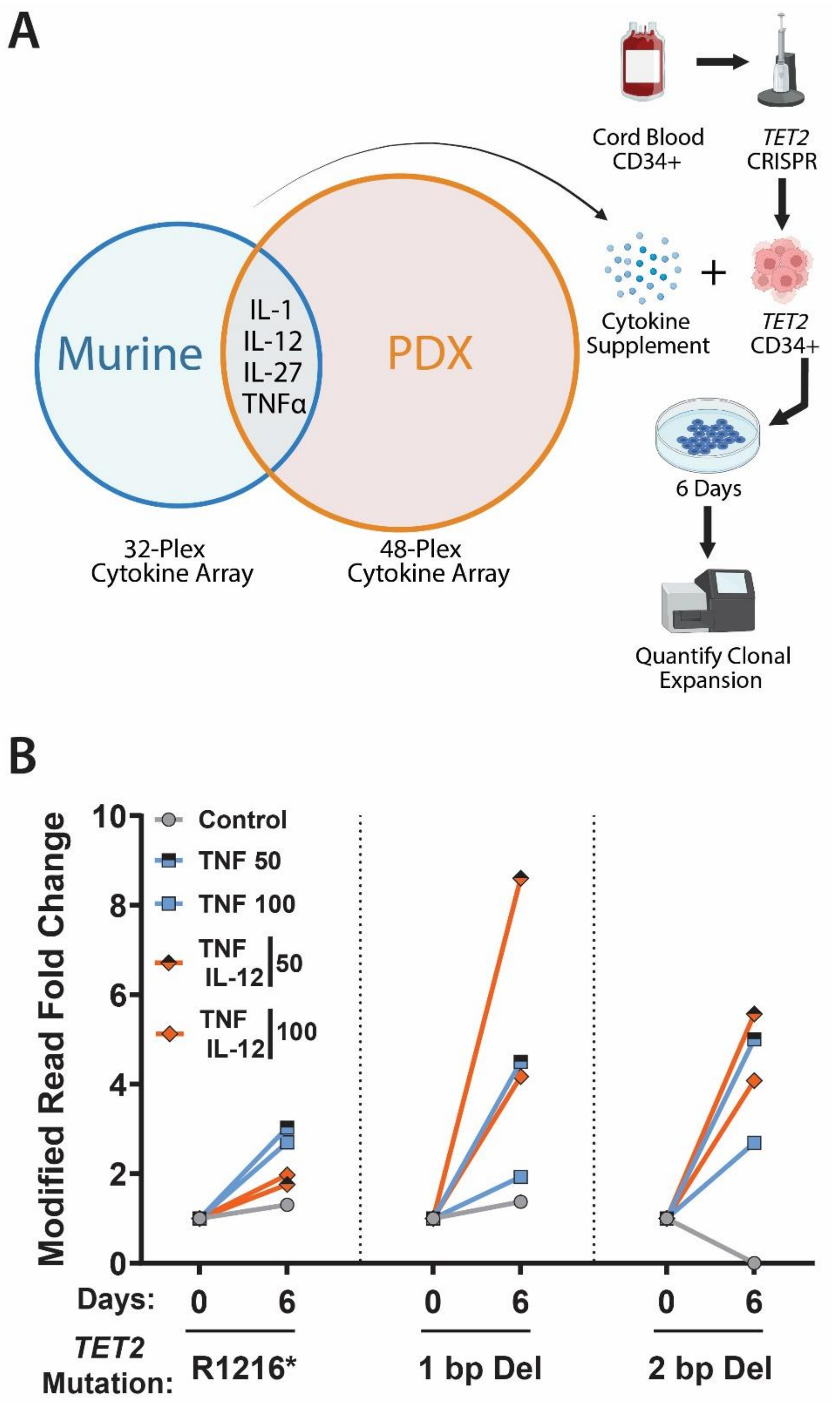


**Fig. S7.**
**(A)** Schematic illustrating workflow for testing cytokines identified to correlate with *TET2*-mutant clone expansion from *in vivo* studies. **(B)** VAF quantification of tracked CRISPR-engineered *TET2* mutations in CB-derived CD34+ cells in media supplemented with indicated cytokines.

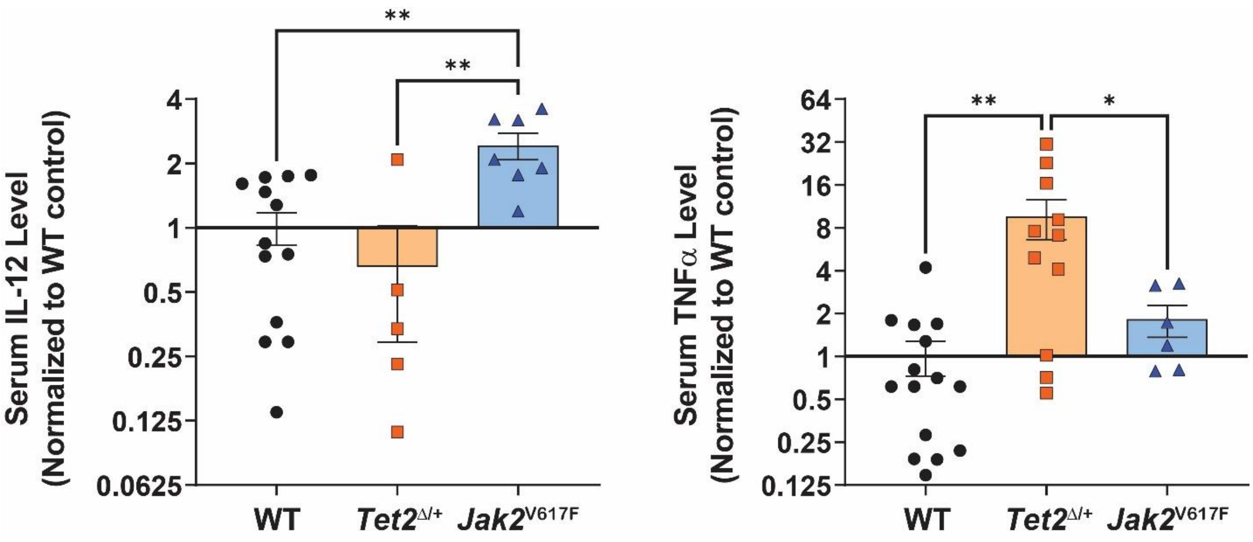


**Fig. S8.**
Serum cytokine levels of IL-12 and TNFα from donor WT, *Tet2*$^{\Delta/+}$ and *Jak2*$^{V617F}$ mice normalized to average levels of WT mice.

| Figure | UPN | Age | Sex | Dx | *JAK2*$^{V617F}$ VAF | Other Variants | Treatments |
|---|---|---|---|---|---|---|---|
| 1 | 950899 | 73 | M | MF | 25% | None Reported | Ruxolitinib + Aspirin |
| 1 | 950899 | 75 | M | sAML | ND | U2AF1, NRAS, KRAS | Transfusion |
| 1 | 374024 | 75 | M | PV | 8% | None Reported | Aspirin |
| 1 | 374024 | 79 | M | sAML | ND | NRAS | Aranesp |
| 1 | 638574 | 59 | M | MF | Positive | None Reported | N/A |
| 1 | 638574 | 60 | M | sAML | ND | RUNX1, PHF6 | N/A |
| 1 | 101811 | 78 | F | PV | 40% | DNMT3A, PPM1D | Hydrea + Aspirin + Intermittent phlebotomy |
| 1 | 867898 | 64 | F | MF | 40% | None Reported | |
| 2 / 4 | 117987 | 38 | M | PV | 5% | None Reported | Aspirin + Intermittent phlebotomy |
| 2 / 4 | 867898 | 64 | F | MF | 40% | None Reported | Momelotinib + Aspirin |
| 2, 4, 5 | 702759 | 26 | F | PV | 2% | None Reported | Aspirin |
| 2 / 4 | 172431 | 76 | F | MF | 86% | BCOR | Ruxolitinib + Retacrit |
| 2 / 4 | 899567 | 58 | F | PV | 88% | None Reported | Hydrea + Aspirin + Intermittent phlebotomy |
| 2 / 4 | 497757 | 55 | F | MF | 47% | None Reported | Ruxolitinib + Aspirin |
| 2 / 4 | 603873 | 49 | M | PV | 5% | None Reported | Aspirin + Intermittent phlebotomy |
| 2 / 4 | 523915 | 73 | M | MF | 10% | ASXL1, CBL, EZH2, SETBP1 | Ruxolitinib |

**Table S1.**

Clinical characteristics from patient samples used throughout the study. All MPN samples contained a *JAK2*$^{V617F}$ driver mutation.

N/A = data not available.

| | | | | |
|---|---|---|---|---|
| *ASXL1* | *ERG* | *KDM6A* | *NRAS* | *SMC1A* |
| *ATM* | *ETV6* | *KIT* | *PHF6* | *SMC3* |
| *BCOR* | *EZH2* | *KMT2A* | *PPM1D* | *STAG2* |
| *BRAF* | *FLT3* | *KRAS* | *PTEN* | *STAT3* |
| *CALR* | *GATA2* | *MPL* | *PTPN11* | *TET2* |
| *CBL* | *GNAS* | *MYC* | *RAD21* | *TP53* |
| *CHEK2* | *IDH1* | *MYD88* | *RUNX1* | *U2AF1* |
| *CSF3R* | *IDH2* | *NF1* | *SETBP1* | *WT1* |
| *DNMT3A* | *JAK2* | *NPM1* | *SF3B1* | *ZRSR2* |

**Table S2.**
MissionBio Tapestri Myeloid Panel targeting 45 genes with 312 amplicons over approximately 65kb of target space.

**Data S1. (separate file)**

Gene expression analysis of 1) *Tet2*$^{\Delta/+}$ HSPCs isolated from a *Jak2*$^{V617F}$ background vs. *Tet2*$^{\Delta/+}$ HSPCs isolated from a WT background, 2) *Tet2*$^{\Delta/+}$ HSPCs isolated from a *Jak2*$^{V617F}$ background vs. WT HSPCs isolated from a *Jak2*$^{V617F}$ background, and 3) WT HSPCs isolated from a *Jak2*$^{V617F}$ background vs. WT HSPCs isolated from a WT background.
